# Engineering ATP Import in Yeast Uncovers a Synthetic Route to Extend Cellular Lifespan

**DOI:** 10.1101/2025.04.14.648769

**Authors:** Naci Oz, Hetian Su, Vedat Sari, Praveen Patnaik, Rohil Hameed, Jong Hee Song, Derek C. Prosser, Vyacheslav M. Labunskyy, Vadim N. Gladyshev, Nan Hao, Alaattin Kaya

## Abstract

Mitochondrial dysfunction and declining ATP production are common features of aging, yet whether ATP availability itself directly regulates cellular lifespan. However, the causal relationship between cellular ATP homeostasis and aging has not been established. Here, we developed a synthetic system to manipulate intracellular ATP independently of endogenous energy production by expressing a plasma membrane–targeted nucleotide transporter, *NTT1*, from the intracellular parasite *Encephalitozoon cuniculi* in *Saccharomyces cerevisiae*. *NTT1* expression depleted intracellular ATP in the absence of extracellular ATP, whereas ATP supplementation produced robust *NTT1*-dependent ATP uptake and increased intracellular ATP abundance. ATP availability strongly influenced replicative lifespan: ATP depletion shortened lifespan, whereas ATP supplementation restored and extended lifespan in NTT1-expressing cells. Unexpectedly, extracellular ATP also extended lifespan in wild-type cells that lack ATP import, revealing an *NTT1*-independent response to extracellular ATP. Transcriptomic analyses showed that *NTT1*- mediated ATP import suppresses glucose uptake, carbohydrate catabolism, mitochondrial respiration, and autophagy, whereas extracellular ATP elicits a distinct transcriptional response in wild-type cells involving metabolic, mitochondrial, and signaling pathways. Single-cell aging analyses further showed that ATP supplementation extends lifespan across distinct aging trajectories and shifts cells away from the mitochondrial dysfunction–associated aging state. Finally, experiments in cells lacking mitochondrial DNA separated these effects mechanistically: *NTT1*-associated lifespan phenotypes and high-ATP toxicity required functional mitochondria, whereas extracellular ATP extended lifespan in wild-type cells independently of mitochondrial respiration. Together, these findings demonstrate that ATP availability is a direct regulator of cellular aging and reveal distinct metabolic and extracellular ATP-responsive routes through which cellular energy state influences longevity.

**Significance:** Cellular energy homeostasis is a crucial factor in determining the health and longevity of organisms. While intracellular ATP levels are tightly regulated, the idea that cells can directly take in extracellular ATP to influence metabolism has not been thoroughly explored. In this study, we engineered yeast cells to import external ATP and demonstrated that this approach significantly alters mitochondrial function, metabolic flow, and aging processes. Our findings show that ATP uptake inhibits catabolic pathways and modulates mitochondrial bioenergetics function, thereby extending cellular lifespan through a novel and non-traditional mechanism. This research reveals an unexpected degree of metabolic flexibility and introduces a synthetic biology-based method to reprogram energy metabolism and longevity. The principles established in this study provide a new framework for understanding the role of cellular bioenergetics in aging, highlighting how the modulation of ATP availability can impact metabolic states and lifespan regulation.

## Introduction

Aging, a complex and multifactorial process, is marked by the gradual decline in physiological function, culminating in increased susceptibility to diseases and mortality (1,2). Central to this process is mitochondrial dysfunction, a critical factor in the cellular and systemic manifestations of aging (3–5). Mitochondria produce ATP through oxidative phosphorylation and regulate various cellular pathways such as calcium homeostasis, redox signaling, and apoptosis (6,7). Over the past decade, a growing body of research has established a strong connection between mitochondrial dysfunction and key characteristics of the aging process. Alterations such as somatic mutations in mitochondrial DNA (mtDNA), impairments in the respiratory chain, and elevated production of mitochondrial reactive oxygen species (ROS) have been implicated. These changes lead to oxidative stress and the accumulation of molecular damage, ultimately disrupting cellular function and promoting senescence and age-related decline. Furthermore, mitochondrial dysfunction, resulting from structural and functional changes, is linked to age-related degenerative diseases (4,8,9). However, recent evidence suggests that mild mitochondrial stress, known as mitohormesis, can promote adaptive responses, enhancing longevity through hormetic mechanisms (5,10–13).

Mitochondrial dynamics, encompassing fusion, fission, and mitophagy, are crucial in maintaining mitochondrial integrity and metabolic and bioenergetic function throughout the lifespan (14–16). As organisms age, disruption in these dynamics can lead to the accumulation of dysfunctional mitochondria. Furthermore, mitochondrial quality control, mediated by mitophagy and fission/fusion processes, declines with age, exacerbating mitochondrial metabolic dysfunction, resulting in an imbalance of energy supply and demand (8,9,14–19). Caloric restriction (CR), a well-established metabolic intervention to extend lifespan, physical activity, and pharmacological interventions targeting conserved longevity pathways improve mitochondrial biogenesis and reduce oxidative damage, indicating that metabolic interventions targeting mitochondria can mitigate age-related imbalance of energy supply and demand (20–23).

The concept of ATP homeostasis, essential for cellular metabolism and signaling, emerges as a pivotal factor in mitochondrial-mediated lifespan regulation (24–26). While several studies have explored how mitochondrial dysfunction impacts aging, few have directly examined the role of intracellular ATP availability and energy supply and demand balance in modulating longevity (6,9,12,24). For example, in *Caenorhabditis elegans*, a mild reduction in mitochondrial metabolic function during developmental stages confers increased lifespan, highlighting the nuanced relationship between energy production and aging (12). Similarly, research on mild mitochondrial uncoupling in mice demonstrates that metabolic shifts favoring increased energy expenditure can reduce ROS production and extend lifespan (11,27).

In this study, we sought to determine whether ATP availability itself directly influences cellular aging. To experimentally manipulate intracellular ATP independently of endogenous ATP-generating pathways, we engineered budding yeast to express a plasma membrane–targeted nucleotide transporter, *NTT1*, from the intracellular parasite *Encephalitozoon cuniculi* (28,29), thereby enabling direct uptake of extracellular ATP. Using this system together with ATP measurements, transcriptomic profiling, and single-cell replicative aging analyses, we found that *NTT1* expression depleted intracellular ATP and shortened lifespan in the absence of extracellular ATP, whereas ATP supplementation increased intracellular ATP and restored and extended lifespan. Elevated ATP through Ntt1-mediated import induced broad metabolic reprogramming, including suppression of glucose utilization, catabolic pathways, mitochondrial respiration, and autophagy. Unexpectedly, extracellular ATP also altered gene expression and extended lifespan in wild-type cells that do not import ATP, revealing a distinct Ntt1-independent cellular response to extracellular ATP.

Analysis of individual aging trajectories further showed that ATP supplementation extended lifespan in both major modes of yeast replicative aging and reduced the proportion of cells entering mode-2 aging, which is associated with mitochondrial dysfunction and ATP decline. Experiments in cells lacking mitochondrial DNA further distinguished the mechanisms underlying these effects. The lifespan defects associated with NTT1 expression and the toxicity caused by high ATP availability in NTT1-expressing cells were suppressed when mitochondrial respiration was eliminated, demonstrating that these effects require functional mitochondria. In contrast, extracellular ATP extended lifespan in wild-type cells even in the absence of mitochondrial respiration. Together, these findings demonstrate that ATP availability directly regulates cellular aging and reveal distinct mitochondria-dependent and mitochondria-independent routes through which ATP influences metabolism, aging trajectory, and replicative lifespan.

## Results

### Overexpression of *NTT1* mediates ATP import into *S. cerevisiae* cells

To alter intracellular ATP levels through extracellular feeding, we created a yeast model expressing a nucleotide transporter (*NTT1*) from the intracellular fungal parasite *Encephalitozoon cuniculi*. *E. cuniculi* naturally uses this exchanger to import ATP from the cytosol of eukaryotic host cells to sustain its survival (28,29). To examine Ntt1 localization in yeast, we modified its sequence by fusing the Pleckstrin homology (PH) domain of *SLM1* to its N-terminus, a well-established plasma membrane targeting signal. We generated expression cassettes with or without a C-terminal GFP tag and integrated them into the genome of wild-type (*Wt-BY4741*) yeast. *NTT1* expression was driven by the constitutive *GPD* promoter (**Fig. 1A**). Fluorescence microscopy confirmed plasma membrane localization of the PH-Ntt1 fusion protein (**Fig. 1B**). This engineered system enables the exchange of extracellular ATP for intracellular ADP via *NTT1* (**Fig. 1C**).

**Figure 1:**
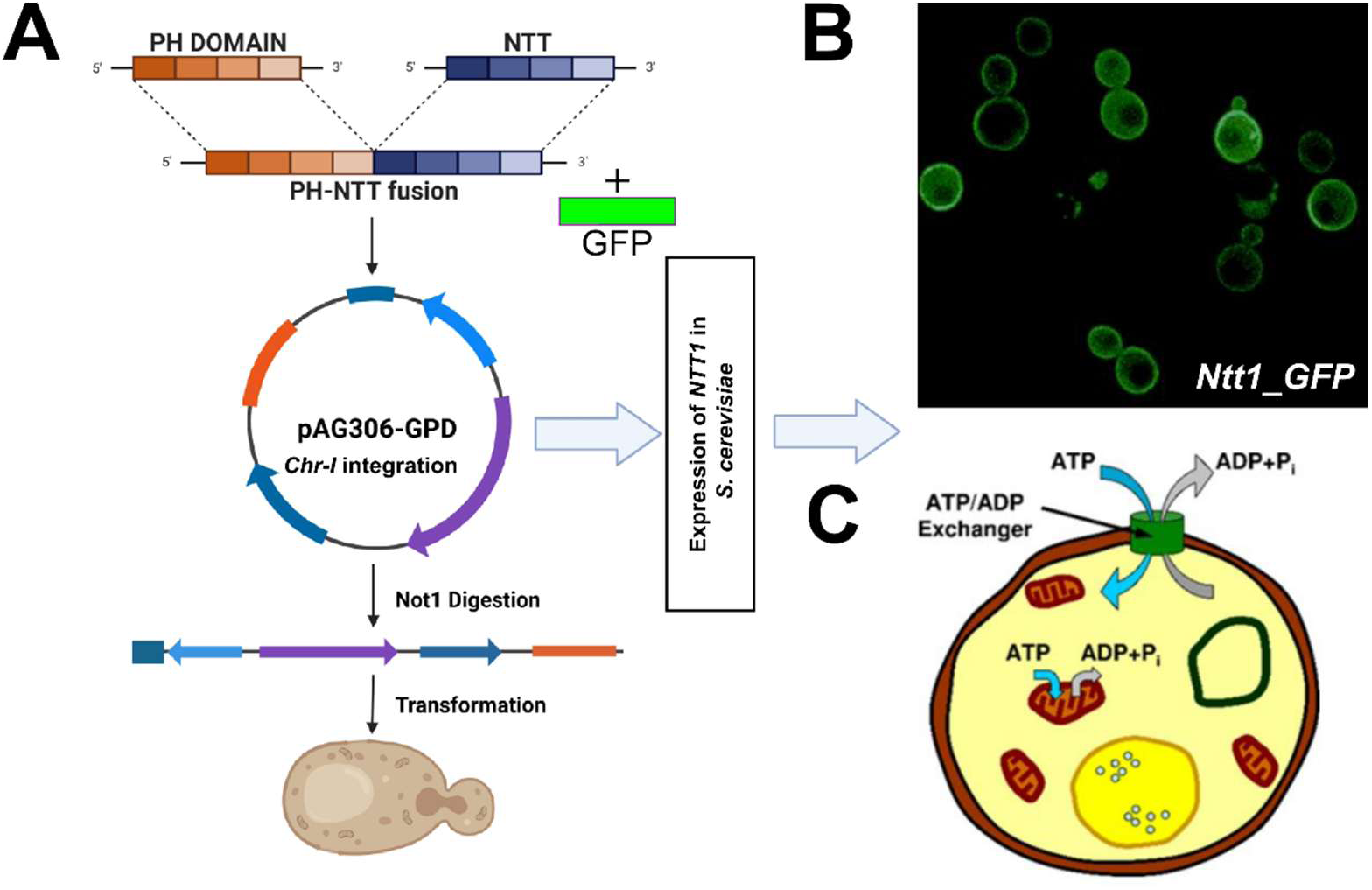
Strain design. (**A**) Pleckstrin homology (PH) domain of *SLM1* gene fused with *NTT1* and inserted in the vector (*pAG306GPD-ccdB-chr1*). The constructed (*PH-NTT1*) plasmid, digested with *NotI*, was integrated into the yeast genome. (**B**) In a separate construct, *yeGFP* sequence was fused to the 3’ end of *PH-NTT1* sequence and expressed in yeast and visualized by confocal microscopy to assess cellular localization. (**C**) Overall yeast strain design with *NTT1* overexpression mediating ATP import from extracellular environment.

Next, to test whether *NTT1* expression mediates ATP import from the extracellular environment, we created a second yeast cell model by integrating the functional *NTT1*-cassette (without GFP) into the genome of a Wt yeast. We also genome-integrated a fluorescent ATP biosensor (Queen), to quantify intracellular ATP levels in single living yeast cells (30,31) (**Fig. 2A**). Based on this reporter strain, our analysis showed that *NTT1* expression in cells severely depleted intracellular ATP levels, compared to Wt control cells (**Fig. 2B**). Supplementing the medium with 5 mM ATP increased ATP significantly (p=7.5E-09), exceeding levels observed in Wt controls, indicating efficient ATP uptake via Ntt1 from the extracellular environment (**Supplementary File 1**). Interestingly, Wt cells lacking *NTT1* exhibited a significant increase in ATP levels (p = 2.22E-16) upon ATP supplementation, though to a lesser extent than NTT1- expressing cells (hereafter referred to as NTT1 cells) (**Fig. 2B**). Since *S. cerevisiae* lacks known plasma-membrane ATP transporters, this suggests that extracellular ATP may activate downstream signaling via unknown nucleotide-sensing receptors, promoting mitochondrial function and ATP synthesis. Alternatively, an unidentified transporter might facilitate ATP uptake.

**Figure 2:**
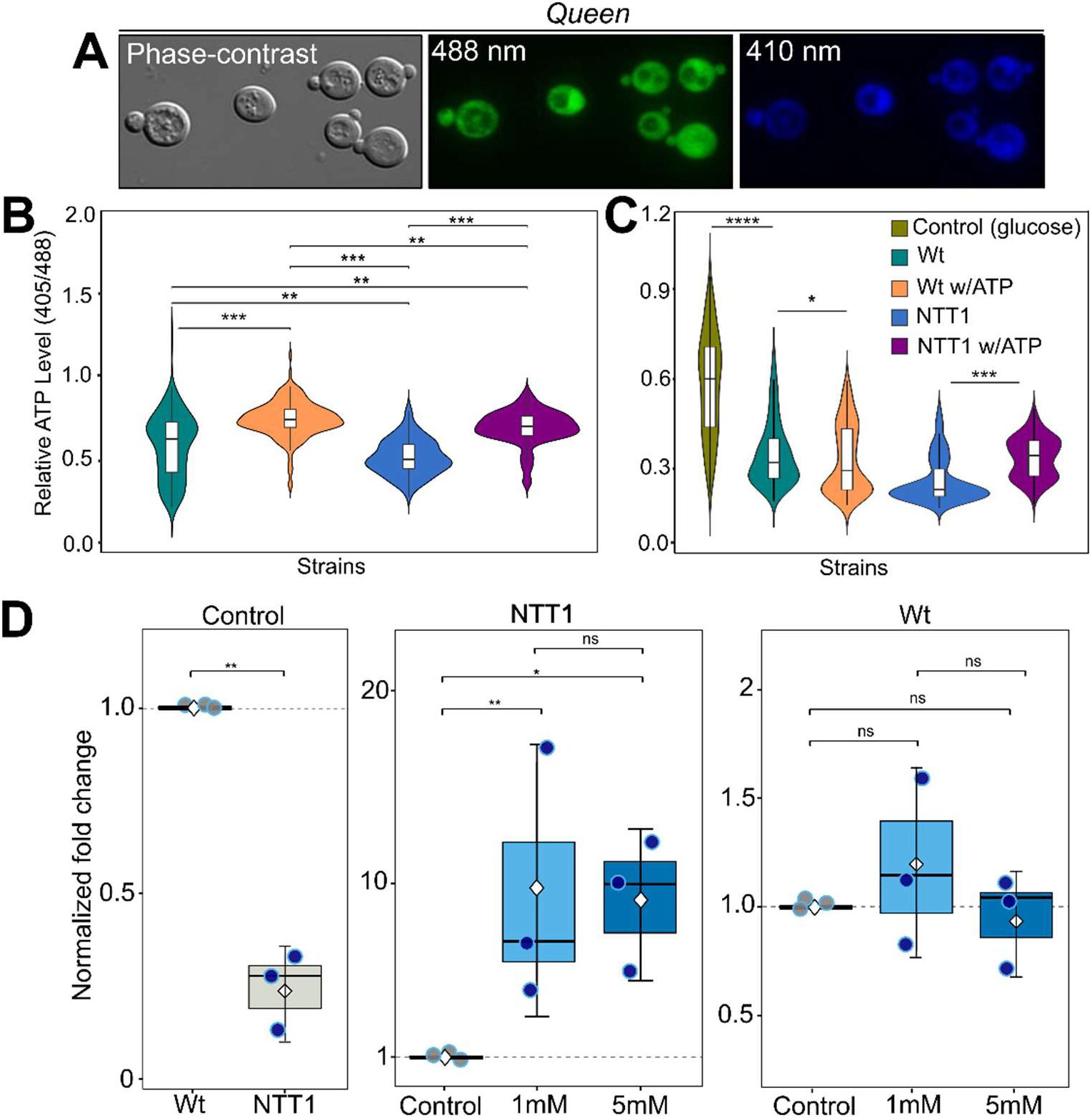
Measurement of relative ATP levels. (**A**) Yeast cells, expressing fluorescent QUEEN ATP reporters were imaged using fluorescence microscopy. Relative ATP levels were measured in cells grown in media containing (**B**) 2% glucose or (**C**) 20 mM 2DG and are presented as violin plots. Sample sizes for each strain are between 265 and 315 cells. Under 2DG condition, ATP treatment did not restore ATP levels in wild-type cells, while NTT1 cells increased intracellular ATP levels (**D**) LC-MS/MS measurement of intracellular ATP changes in Wt and NTT1 expressing cells. A non-parametric Kruskal-Wallis test was used for comparison of groups for the QUEEN measurements. For the LC-MS/MS data pairwise t-test was used to compare the means of treated versus non-treated samples of each strain and determine if a statistically significant difference exists between them. Statistical significance between groups and raw QUEEN ratio as well as LC- MS/MS data are available in **Supplementary File 1.**

To validate these findings, we designed an experiment to inhibit endogenous ATP synthesis by treating cells with 2-deoxyglucose (2DG), a glucose analog that competitively inhibits glycolysis. Once phosphorylated by hexokinase, 2DG becomes 2DG-6-phosphate, which accumulates and inhibits glycolysis, leading to rapid ATP depletion (32). As expected, 30 minutes of 2DG treatment sharply reduced intracellular ATP levels in both wild-type and NTT1 cells (**Fig. 2C**). When NTT1 cells were subsequently treated with 5 mM ATP, intracellular ATP levels increased, confirming ATP import via *NTT1*. In contrast, ATP treatment did not restore ATP levels in wild-type cells, arguing against the presence of an unidentified ATP transporter (**Fig. 2C and Supplementary File 1**).

Finally, we further validated the *NTT1*-mediated ATP import by analyzing cells with LC-MS/MS after ATP treatments. Consistent with the QUEEN measurement of control cells, our analysis revealed significantly decreased ATP level in *NTT1* cells under non-treated condition in comparison to the Wt controls (p=0.008) (**Fig. 2D**). Similarly, 30 minutes treatment of 1mM and 5mM exogenous ATP caused 6- and 10-fold increase in intracellular ATP in *NTT1* expressing cells respectively (p < 0.05). For the Wt cells, LC-MS/MS analyses revealed no significant (**Fig. 2D and Supplementary File 1**) alteration in intracellular ATP level under both treatment conditions (1mM and 5mM). The two methods (QUEEN versus LC-MS/MS) may have different sensitivities and linear ranges that might cause these discrepancies, observed for the Wt cells. However, these data further validate our observation that Wt cells do not import exogenous ATP (**Fig. 2C**). The QUEEN reporter provided real-time, dynamic measurements of ATP concentration in living cells under continuous treatment of freshly supplemented medium conditions, mediated by NTT1 import mechanism. This allows it to capture rapid, transient increases or fluxes in ATP that occur immediately after the continuous exogenous treatment. However, since there is no ATP import in Wt cells, ATP treatment through continuously supplemented medium might be needed to induce downstream signaling to induce ATP synthesis. Furthermore, LC-MS/MS typically involves sample extraction and processing steps that effectively provide a "snapshot" of the average, steady-state ATP levels across the cell population at a single point in time. The transient slow alteration of ATP abundance detected by QUEEN in Wt cells may be rapidly metabolized or pumped out of the Wt cells before the LC-MS/MS sample preparation. Considering the rapid import and increased abundance of ATP in *NTT1* cells, which is detected by using both Queen and LC-MS/MS, Wt cells might need longer and continuous treatment conditions. Overall, these data further validate modulation of intracellular ATP levels through extracellular feeding in cells harboring functional *NTT1*.

### Altered ATP level targets cellular metabolic pathways

To investigate the effect of exogenous ATP treatment on gene expression in Wt and NTT1 cells, we performed RNA-seq analysis under both treated and non-treated conditions (**Fig. 3A**, **Supplementary File 2**). Our goal was to distinguish molecular signatures triggered by extracellular ATP-treatment in control cells from those driven by increased intracellular ATP via Ntt1-mediated uptake in engineered cells. First, to define the host response to expression of *NTT1* in the absence of exogenous nucleotide, we performed RNA-seq on yeast cells carrying a chromosomally integrated *NTT1* cassette against an isogenic Wt control. Differential expression analyses revealed that at a nominal threshold of p < 0.05, 360 genes were differentially expressed (DEGs) between NTT1 and Wt cells, of which 22 passed FDR correction (Benjamini–Hochberg-adjusted p < 0.05). However, further analyses of these DEGs revealed that the response was strikingly asymmetric: 268 genes (74 %) were down-regulated and only 92 (26 %) were up- regulated, a bias significantly different from chance (binomial p = 4.5 × 10^−21^; **Fig. S1A, B**). This skew indicates that NTT1 expression does not simply rewire host transcription in both directions but predominantly attenuates expression of a defined set of biosynthetic and transport functions, with a much smaller induction program at gene expression level.

**Figure 3:**
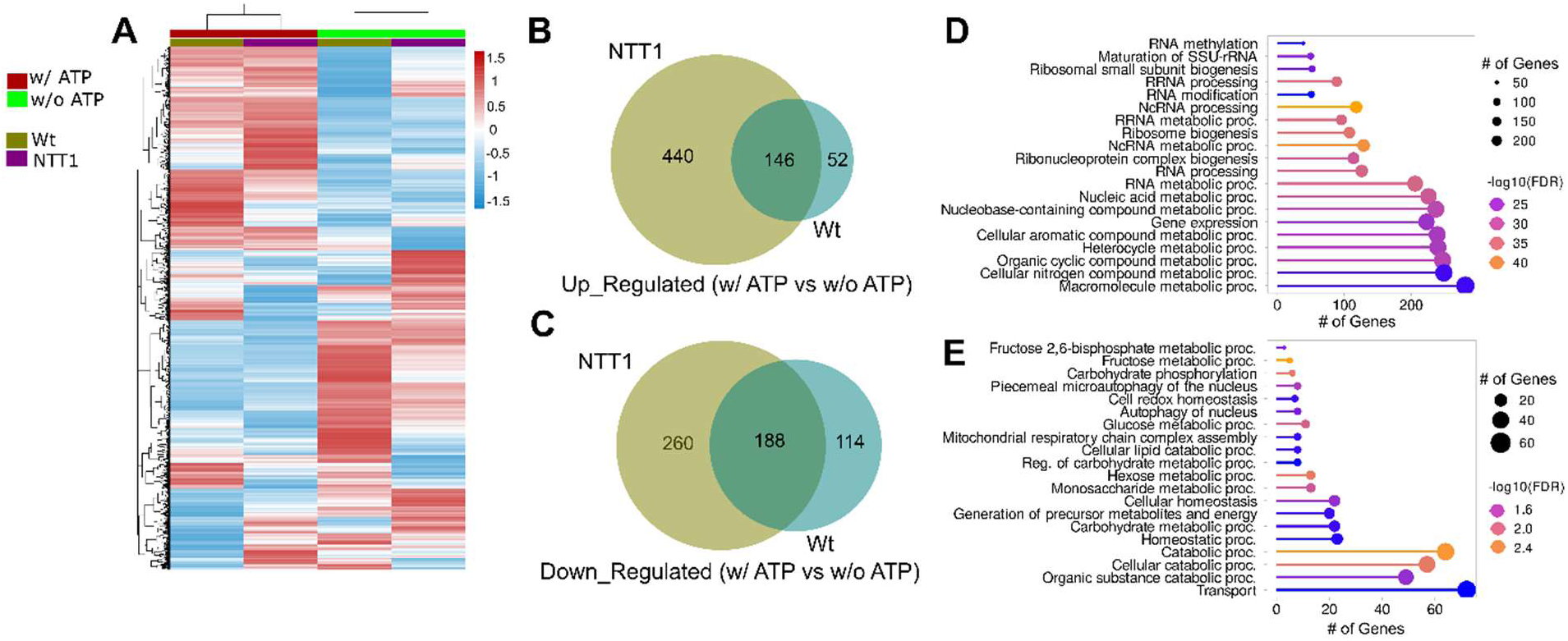
Transcriptomic analysis of Wt and NTT1 cells with and without ATP treatment. (**A**) Heatmap showing hierarchical clustering of gene expression profiles from Wt and NTT1 cells cultured with ATP (red) and without ATP (green). Expression values are Z-score normalized, with red indicating higher expression and blue indicating lower expression. Three independent cultures for each group were used to isolate total RNA and perform RNA-seq analysis and each cell in the columns shows the mean value from three biological replicates. (**B, C**) Venn diagrams displaying the number of genes uniquely or commonly (**B**) upregulated or (**C**) downregulated in Wt and NTT1 cells treated with ATP compared to untreated controls. (**D, E**) Enrichment analysis showing significantly enriched gene ontology (GO) terms associated with genes (**D**) upregulated and (**E**) downregulated in NTT1 cells compared to their corresponding untreated controls. Circle size reflects the number of genes in each GO term, and color indicates significance level as represented by -log10 (FDR). The normalized counts and the complete list of DEGs with *p* values can be found in **Supplementary File 2**.

The strongest functional enrichment among significantly up-regulated genes was for the mitochondrial inner-membrane biogenesis machinery (adjusted p = 1.1×10^−5^; **Fig. S1C**). This included components of the mitochondrial ribosome (*MRPL4*, *MRPL10*, *MRPL16*, *MRPL23*, *MRPL32*, *MRPL33*, *MRPL35*, *MRPL37*), cytochrome *c* oxidase assembly factors (*COX17*). Along with the nine other OXPHOS-related gene set enrichment (adjusted p = 0.049), the signature is consistent with a *HAP4*-mediated respiratory induction (33) and suggests that cells coordinately raise mitochondrial inner-membrane assembly capacity to compensate for the decreased intracellular ATP level in cells with *NTT1* overexpression (**Fig. S1C**).

On the other hand, the most significant functional enrichment (adjusted p = 6.0×10^−11^) for the down regulated genes was coordinated down-regulation of de novo purine biosynthesis and one-carbon/folate metabolism of the canonical BAS1/PHO2 adenine regulon (**Fig. S1C**), including the entire IMP-biosynthetic arm (*ADE1*, *ADE3*, *ADE5,7*, *ADE8*, *ADE12*, *ADE13*, *ADE17*), the BAS1- co-regulated histidine branch (*HIS4*), and the one-carbon supply genes that feed N10-formyl-THF into purine ring synthesis (*ADE3*, *MTD1*, *SHM2*, *GCV2*) and the NADP kinases (*POS5, UTR1*) (**Fig. S2A**). This data suggests that *NTT1* expression specifically resulted in decreased AMP and downstream ADP synthesis. To validate this observation, we analyzed both ADP and AMP levels in Wt and NTT1 cells. Our analyses revealed that *NTT1* expression resulted in decreased AMP and ADP levels (**Fig. S2B and Supplementary File 1**). Significant (p < 0.05) down regulation of six genes (*KOG1, MEP1, TOR1, TOR2, TCO89, GLN3*) also resulted in enrichment for TOR/Nutrient sensing pathway (adjusted p = 1.2×10^−3^), consistent with an energy crisis caused by NTT1 overexpression (**Fig. S1C**).

Next, we analyzed differentially expressed genes (DEGs) under ATP-treated and untreated conditions for both strains and compared their expression profiles. Extracellular ATP elicited a substantial transcriptional response in wild-type cells (500 DEGs; 198 up, 302 down; **Fig. 3B, C**) despite the fact that yeast cannot import nucleotides across its plasma membrane, establishing that *S. cerevisiae* possesses an active extracellular ATP-sensing mechanism. NTT1 expression approximately doubled the response to ATP treatment (1,034 DEGs; 586 up, 448 down; **Fig. 3B, C**) and inverted the up:down ratio from 0.66 to 1.31. Among these, 146 upregulated and 188 downregulated genes were shared between the two strains, suggesting that these changes result directly from ATP treatment and occur independently of *NTT1* expression (**Fig. 3B, C**).

The dominant induction in common datasets was *RGS2* (log_2_FC = +1.74 in Wt, +1.88 in NTT1+ATP; padj < 10^−15^ in both), the glucose-mediated negative regulator of G-protein signaling that limits Gpa2 output to adenylyl cyclase and whose induction is the canonical feedback marker of an active Gpr1–Gpa2–cAMP–PKA module (34). Further, decreased expression of GPA2 and PKA subunits TPK1 and TPK2 as well as increased expression of PDE2 may reflect compensatory feedback that dampens hyperactive PKA signaling. Consistent with PKA activation, transcripts encoding PKA-repressed factors *YAK1*, *GIS1*, and *MSN4* were reduced (**Fig. S3**). Together, these data implicate the Gpa2–cAMP–PKA pathway as a GPCR-coupled route through which yeast respond to extracellular ATP.

To further dissect ATP-mediated responses in Wt cells, we performed Gene Ontology (GO) (**Fig. S4A, B**) and KEGG pathway enrichment analyses (**Fig. S4C, D**). ATP treatment led to significant upregulation of genes involved in ribosome biogenesis, ATP biosynthesis, MAP kinase (MAPK) signaling, and oxidative phosphorylation (**Fig. S4A, C**). Conversely, downregulated genes were enriched in carbohydrate metabolism and uptake, protein phosphorylation, autophagy, and meiosis pathways (**Fig. S4B, D**). These findings indicate that extracellular ATP triggers intracellular signaling cascades capable of modulating mitochondrial function, cytosolic translation, and protein phosphorylation (**Supplementary File S2**). Notably, genes in the MAPK cascade were significantly enriched (**Fig. S5**), suggesting this pathway might play a key role in mediating the downstream ATP sensing response in Wt cells.

Next, we applied the same enrichment analyses for the DEGs identified by comparing NTT1 cells (treated vs non-treated). We found a complete rewiring of transcriptional regulation of metabolic genes in NTT1 cells following ATP treatment. Like Wt, NTT1 cells also showed upregulation of ribosome biogenesis and RNA processing genes (**Fig. 3D and Fig. S6**). However, while ATP treatment upregulated genes associated with mitochondrial respiratory activity, oxidative phosphorylation, and MAPK signaling in Wt cells (**Fig. S4**), genes from these pathways were downregulated in *NTT1* cells (**Fig. 3E and Fig. S6**).

Further, the downregulated gene set in NTT1 cells was enriched in glucose uptake, energy generation from precursor metabolites, carbohydrate phosphorylation, general catabolic processes, and autophagy (**Fig. 3E and Fig. S6**). This repression mirrors Snf1-mediated metabolic rewiring. *SNF1*, the yeast homolog of mammalian AMP-activated protein kinase (AMPK), plays a pivotal role in energy homeostasis by modulating metabolic responses and glucose de-repression under varying nutrients and ATP levels (32). Low Snf1 activity is expected under high ATP concentrations (35).

To confirm this, we examined the expression of *SNF1* downstream targets associated with nutrient stress. We observed significant downregulation of several genes, including *MIG1*, *TDA1*, *HXK1*, *HXK2*, and *HXT3,* which are involved in glucose transport and metabolism (**Fig. S7; Supplementary File S2**). Additionally, *RGT1*, a major transcription factor that represses glucose transporter (*HXT*) genes under high-glucose conditions, was significantly upregulated in *NTT1* cells (**Supplementary File S2**). Together, these results indicate repression of glucose uptake and metabolism, further supporting the presence of a global metabolic rewiring in response to ATP elevation in NTT1 cells (36,37).

Collectively, our data revealed a striking biological phenomenon: *NTT1*-mediated ATP import suppresses glucose uptake and catabolism, dampening mitochondrial energy production and reconfiguring cellular metabolism. Given that catabolic pathways are a major source of intracellular damage through the production of reactive by-products, this reduction in metabolic flux may also minimize damage accumulation. Consequently, autophagy, a key mechanism for clearing damaged macromolecules, was downregulated in these cells. These findings highlight two mechanistically distinct but complementary models by which ATP influences cellular physiology and lifespan. In NTT1 cells, ATP is directly imported from the extracellular environment, leading to a robust and specific rewiring of cellular metabolism characterized by the suppression of glucose uptake, glycolysis, and mitochondrial respiration. This *NTT1*-dependent ATP influx mimics a high-energy state, repressing energy-generating catabolic pathways and reducing the need for damage-clearing processes such as autophagy, potentially preserving cellular integrity over time. In contrast, extracellular ATP also exerted effects in Wt cells lacking *NTT1*, suggesting a separate, *NTT1*-independent mechanism possibly mediated through extracellular ATP sensing. Based on altered gene expression from the transcriptome analyses, our data suggest that Gpa2–cAMP–PKA pathway and MAPK cascade might mediate the sensing of extracellular ATP, that cells can detect and respond to extracellular nucleotides without direct ATP uptake.

In yeast, multiple MAPK pathways branch off and share components to regulate various cellular responses. These responses include pheromone signaling, filamentous growth (also known as pseudohyphal or starvation-induced growth), the high osmolarity glycerol (HOG) pathway, and other stress response pathways (38–40) The pheromone response, and filamentous growth pathways are both initiated by G-protein signaling and share common components such as the MAPKKK Ste11p and MAPKK Ste7p. In contrast, the HOG pathway comprises two parallel branches (Sln1 and Sho1) that converge at a common MAPKK (Pbs2) and MAPK (Hog1). (**Fig. S5**). Our data indicated that the expression pattern of gene components of G-protein signaling cascades might be regulated positively while the Hog1 mediated osmolarity pathway was repressed (**Fig. S5**). To further investigate whether Hog1 activity is altered when cells are treated with ATP, we performed western blot analyses to analyze the expression and phosphorylation level of Hog1 (Hog1^p^) protein with two independent antibodies from different sources (please see method) developed specifically against Hog1^p^ (**Fig. 4A,B and Fig. S8**). We found that ATP treatment decreased the Hog1^p^ level in both Wt and NTT1 cells, consistent with reduced Hog1 activity and repression of the HOG pathway. To investigate whether reduced Hog1 activity is dependent on mitochondrial function, we further investigated Hog1^p^ levels in *rho^0^* isolates of both cell types. The HOG1-pathway both promotes and regulates mitochondrial function and selective mitochondrial degradation (39,41). Prolonged activation was also associated with mitochondria mediated increase in reactive oxygen species (ROS) levels (38,41). Although, in Wt *rho^0^* cells Hog1^p^ level decreased, we did not observe overall change in Hog1^p^ level under ATP treatment conditions in NTT1 cells. (**Fig. 4A-C**). These data suggest that the presence and absence of respiratory active mitochondria can mediate signaling cascades differentially, when cells are treated with extracellular ATP in NTT1 but not in Wt cells. Furthermore, it was previously reported that activation of PKA suppresses the phosphorylation of Hog1, and downstream of the HOG pathway, aligning with our transcriptome data suggesting that ATP treatment activates PKA pathway and this activation might be associated with decreased Hog1 activity (42,43) in Wt cells independent of mitochondrial activity, however our data suggests that PKA-mediated repression of HOG pathway might be dependent on presence respiratory active mitochondria in NTT1 cells.

**Figure 4.**
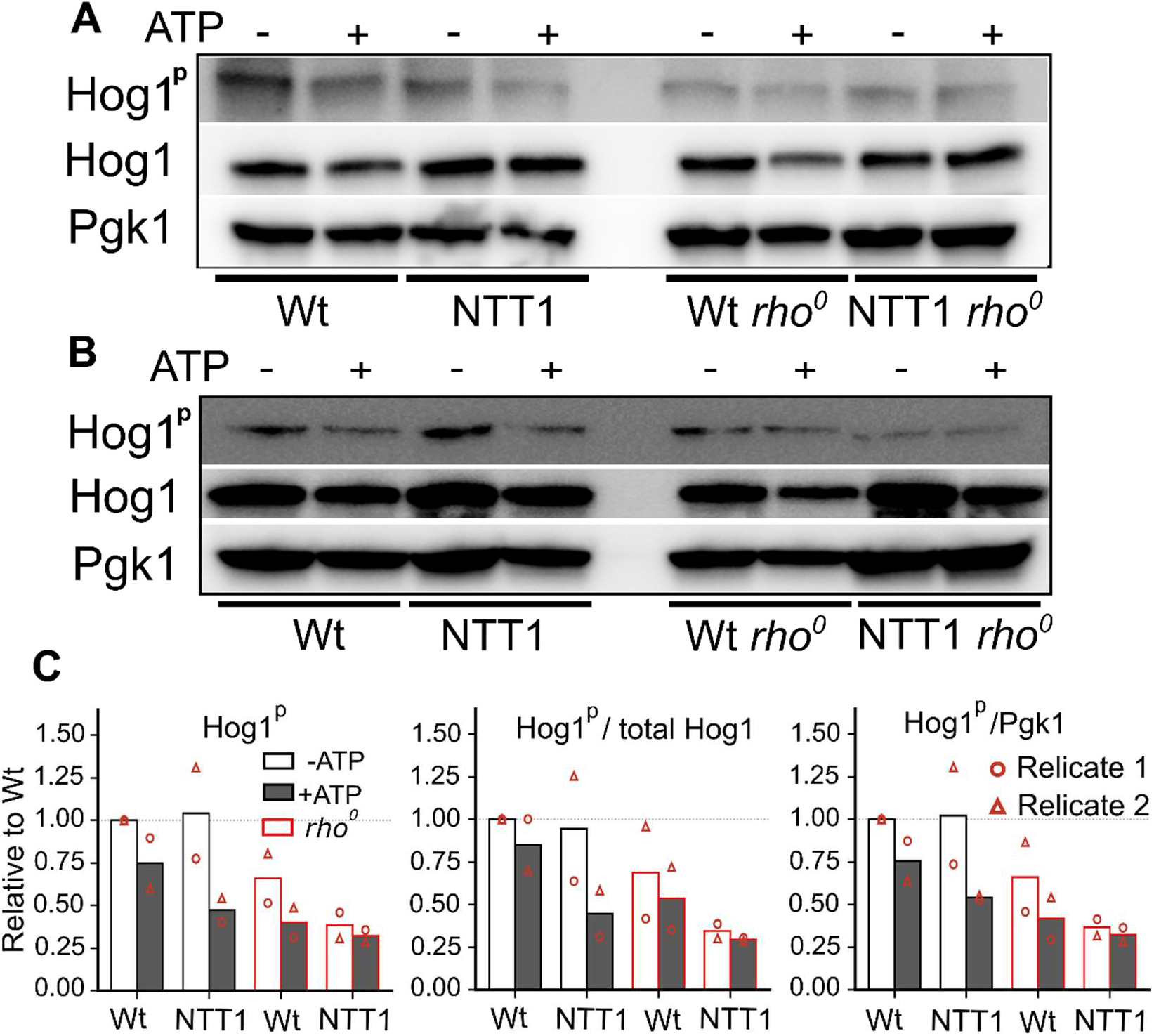
ATP treatment represses Hog1 activation through a mitochondria-dependent mechanism in NTT1 but not in Wt cells. **(A and B)** The effect of 5 mM ATP treatment on Hog1 activation was assessed by Western blot analysis using two independent antibodies against phospho-Hog1 in two independent biological replicates (see Method). Pgk1 was used as an internal loading control. Uncropped membranes are provided in Fig. S8. (**C**) Densitometric quantification of Hog1 phosphorylation. Quantification of the immunoblots in Figure 4A, B. Hog1^P^ signal is shown as background-subtracted integrated density (left) and normalized to total Hog1 (center) or to Pgk1 (right) in the same lane, each expressed relative to untreated wild-type cells. Open circles and triangles show the individual replicate values and red bars represent the *rho^0^* isolates of each cell types.

Together, these data from two cell types (Wt versus NTT1 cells) suggest that ATP can modulate cellular pathways through intracellular energy repletion, extracellular ATP sensing, and activating downstream signaling, revealing a compartment-specific role of ATP. The unexpected divergence of ATP treatment between NTT1 cells and Wt underscores the complexity of regulating cellular energy demand, further validating ATP as both a metabolic substrate and a signaling molecule with profound implications in cellular homeostasis.

### Altered intracellular ATP level modulates lifespan

We further investigated the consequences of altered ATP abundance on cellular aging. Due to the unstable nature of ATP molecules in the culture medium, we employed a state-of-the-art microfluidic aging chamber (44–48), which allowed cells to be continuously exposed to a medium supplemented with fresh ATP. This system was also integrated with high-resolution fluorescence microscopy, enabling real-time measurement of ATP levels throughout the yeast lifespan at the single-cell level using the fluorescence intensity of a novel and reliable Queen-ATP reporter (30,31). Although the QUEEN reporter measures ATP level by the ratio of fluorescence intensities at 410 nm and 488 nm excitation wavelengths, repeated and prolonged exposure to high-intensity 410 nm excitation causes phototoxicity in yeast cells during long-term aging experiments. Therefore, only 488 nm excitation was used in these experiments, and the resulting QUEEN fluorescence signal is inversely correlated with intracellular ATP levels (**Fig. 5A-C**). Our results revealed that Wt cells exhibit divergent ATP dynamics during aging. While some cells showed increased ATP levels, others maintained unchanged or reduced ATP levels (**Fig. 5A**), consistent with previous reports describing two modes of yeast aging, linked to mitochondrial function and *SIR2* activity (44,48–51). NTT1 cells displayed a similar divergence but with greater variability (**Fig. 5B**), and a shortened lifespan (**Fig. 5D**). We next assessed age-associated ATP dynamics and lifespan under ATP supplementation. Wt cells exhibited a significant increase in ATP levels (**Fig. 5A, C**) along with an extended lifespan (**Fig. 5D**) upon 10 µM ATP supplementation. NTT1 cells showed an even more pronounced increase in ATP levels (**Fig. 5B, C**), accompanied by a significant extension (p = 4.03E-18) in lifespan (**Fig. 5D**).

**Figure 5:**
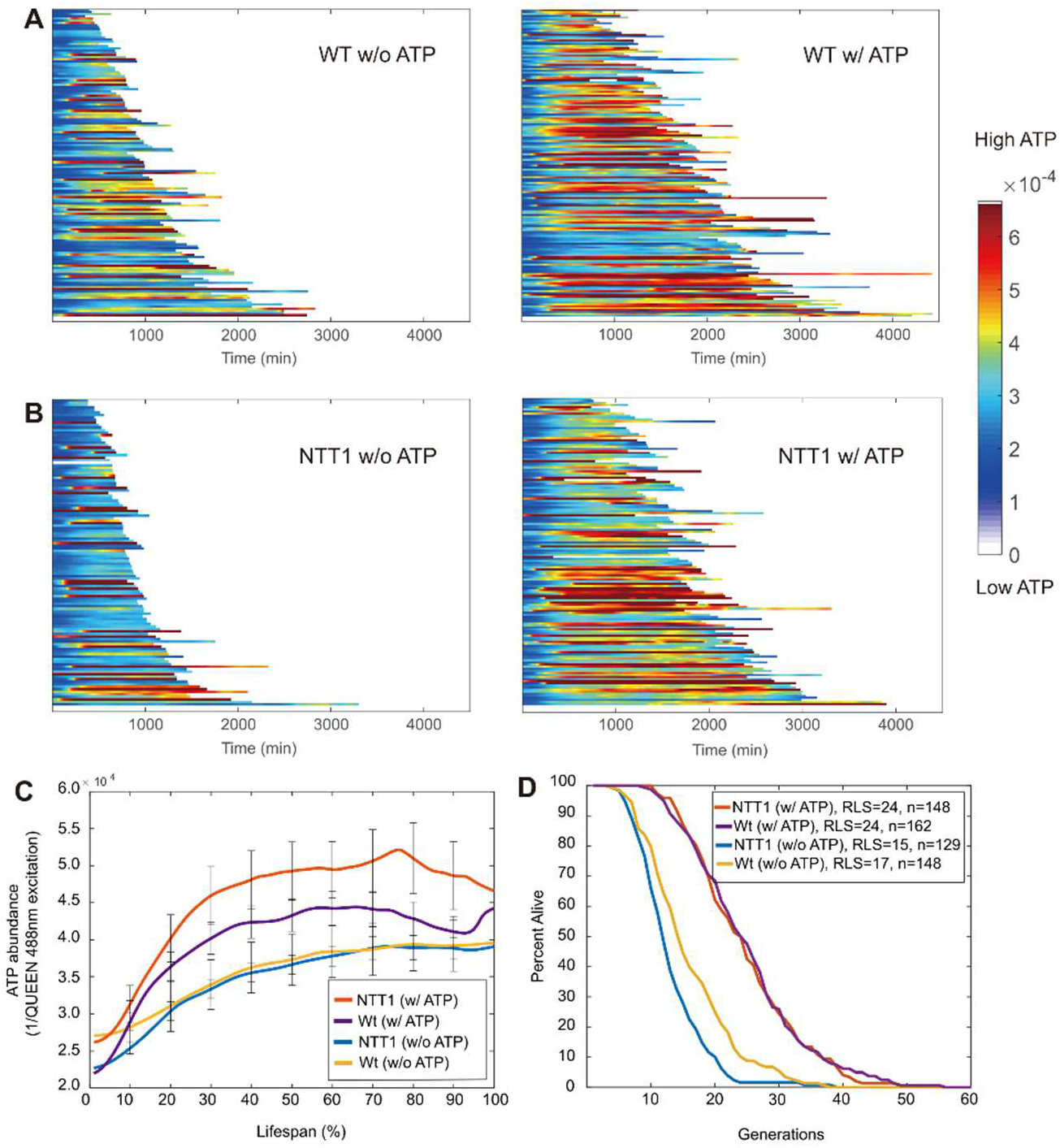
Quantifications of ATP levels and replicative lifespans of the Wt strain and NTT1 strain under ATP treatment conditions. Heatmaps show intracellular ATP abundance in (**A**) WT and (**B**) NTT1 cells, as measured by the inverse of QUEEN fluorescence at the 488nm excitation (see Method), throughout entire lifespans of single cells. Each row represents a color-coded time trace of a single yeast cell. Left - no ATP supplementation; Right - 10 µM ATP supplementation. Color bar represents the relative ATP abundance (e.g., red color indicates higher ATP) (**C**) Average of single-cell time traces of ATP dynamics aligned by percentage of lifespan. Error bars are standard errors. (**D**) Replicative lifespans of Wt and NTT1 cells, with or without ATP supplementation. Wt: w/ ATP vs w/o ATP (p = 1.28E-12); NTT1-expressing cells: w/ ATP vs w/o ATP (p = 4.03E-18). P-values were calculated with Mann-Whitney U test. The raw fluorescence measurements and lifespan data and number of cells analyzed in each experiments can be found in **Supplementary File 3.**

Together, these findings indicate that intracellular ATP dynamics during aging are heterogeneous and closely linked to lifespan outcomes. Enhancing ATP availability through *NTT1* expression promotes longevity, suggesting that maintaining cellular energy homeostasis is a key determinant of aging trajectories in yeast. In addition, the Wt results are consistent with our transcriptomic data, implying that extracellular ATP may activate downstream signaling through nucleotide-sensing receptors, thereby increasing mitochondrial function, and transient ATP production, and other canonical pathways.

### ATP homeostasis dictates aging pathway selection and longevity in yeast

To further understand how ATP availability influences aging dynamics, we examined the two known trajectories of yeast replicative aging mode-1 and mode-2 under both ATP-treated and untreated conditions (44). These modes are distinguished by their daughter cell morphology: mode-1 cells produce elongated daughters, whereas mode-2 cells give rise to small, round daughters. At the molecular level, mode-1 aging is characterized by rDNA instability and nucleolar enlargement and fragmentation, yet mitochondria remain functional. In contrast, mode-2 cells exhibit impaired mitochondrial function (44,48–51). Consistent with a previous report (40), we found that approximately 60% of Wt cells aged through mode-1, while 40% aged through mode-2 under control (no ATP treatment) conditions. Cells aged through mode-1 exhibited significantly longer lifespans than mode-2 cells (**Fig. 6A**).

**Figure 6:**
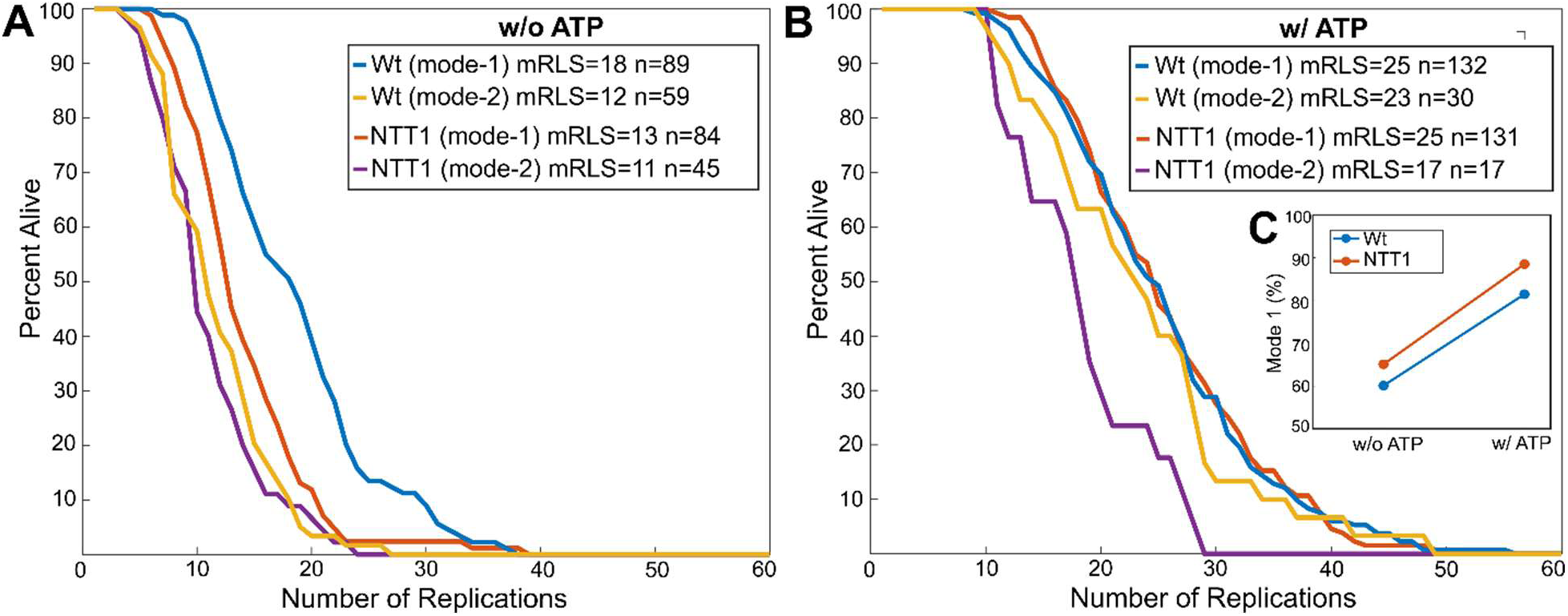
Replicative lifespan of different aging modes and mode distribution under ATP treatment conditions. (**A**) Without ATP, the mode-2 lifespans between the Wt and NTT1 cells showed no significant difference (p = 0.74). On the other hand, NTT1 cells showed significantly shorter mode-1 lifespan in comparison to the Wt control (p = 2.39E-4). (**B**) With ATP, the NTT1 strain showed significantly increased mode-1 lifespan (p = 1.24E-16), but not mode-2 lifespan (p = 0.15), whereas the Wt strain showed significantly increased mode-1 lifespan (p = 3.17E-7) and also mode-2 lifespan (p = 5.98E-6), which was similar to mode-1 lifespan (p = 0.4). The mode-1 lifespans between the 2 strains were similar (p = 0.849). Figures indicate mean replicative lifespan (mRLS) values and number of individual mothers tested in each assay (n) were also included (**C**) Both strains show increased percentage of mode-1 cells under the effects of ATP. The p-values were calculated using Mann-Whitney U test. The lifespan data can be found in **Supplementary File 3.**

We previously showed that mode-1 cells can maintain a relatively constant intracellular ATP level during aging, whereas mode-2 cells show a rapid depletion of ATP due to mitochondrial dysfunction (50). Therefore, Ntt1-mediated ATP depletion may have a more dramatic effect on the lifespan of mode-1 cells than that of ATP-depleted mode-2 cells. As expected, under the condition of no ATP treatment, *NTT1* expression caused a significant reduction in the lifespan of mode-1 cells (Wt mode-1, mean RLS = 18 vs. *NTT1* mode-1, mean RLS = 13; p = 2.39E-4), but not in mode-2 cells (Wt mode-2, mean RLS = 12 vs. *NTT1* mode-2, mean RLS = 11, p = 0.74) (**Fig. 6A**). We then analyzed the effect of ATP treatment. In both Wt and NTT1 cells, ATP treatment increased the lifespans of mode-1 and mode-2 cells (**Fig. 6B**). Furthermore, the treatment also enhanced the proportion of mode-1 cells (**Fig. 6C**), suggesting that ATP treatment can compensate for the age-induced ATP decline in mode-2 cells, biasing the fate decision toward mode-1 aging.

These results indicate that elevating intracellular ATP, whether through *NTT1*-mediated import or via extracellular sensing, can extend cellular lifespan and influence the trajectory of aging. By restoring energy balance, ATP not only delays senescence but also shifts cells toward aging modes associated with greater mitochondrial integrity and extended replicative potential.

### *NTT1*-driven ATP import alters lifespan through mitochondria-dependent mechanisms

To further understand the downstream effect of altered extracellular and intracellular ATP abundance and whether it interferes with mitochondrial function, we designed experiments with *rho^0^* isolates, in which we eliminated mitochondrial DNA (mtDNA). These cells are unable to perform oxidative phosphorylation (OXPHOS) because they lack the necessary components of the electron transport chain encoded by the mitochondrial genome (52,53). RLS of Wt and NTT1 cells, as well as their *rho^0^*isolates, were assayed under the control and higher dose (10 mM) ATP-supplemented medium conditions using a microfluidic aging chamber.

A 10 mM ATP treatment caused a ∼30% mean increase (p =4.52E-12) in the replicative lifespan (RLS) of Wt cells (**Fig. 7A**). This result is similar to the lifespan change seen with a 10 µM treatment (**Fig. 5D**). However, for the *NTT1* cells, in comparison to the 10 µM treatment condition, in which *NTT1* cells extended lifespan to the level of treated Wt cells (**Fig. 5D**), the positive effect of ATP treatment on lifespan was strongly attenuated relative to WT, consistent with dose-dependent toxicity (p= 1.25E-05, Wt treated versus, NTT1 cells treated) (**Fig. 7A**). However, we made an interesting observation that the toxicity effect of 10 mM ATP treatment on lifespan of NTT1 cells was alleviated in *NTT1 rho0* isolates. The 10 mM ATP treatment significantly (p=3.07E-05) increased the lifespan of *NTT1 rho^0^* cells (**Fig. 7B**) comparable to the Wt cells, further supporting the conclusion that mitochondrial function mediates ATP toxicity. In addition, eliminating mtDNA prevented the adverse lifespan effect (**Fig. 5D and Fig. 7A**, **Supplementary File 3**) of *NTT1*-expression, consequently *NTT1 rho^0^* cells displayed lifespan restored to the level of Wt cells (**Fig. 7B**) under the no treatment condition. It should be noted here that under the same condition, we did not observe any significant lifespan differences between Wt and Wt *rho^0^* cells (**Fig. 7A, B**), further confirming that eliminating mtDNA was the primary driver of the increased lifespan for NTT1 cells under both treatment and non-treatment conditions. These data strongly suggest that the toxicity associated with *NTT1* overexpression and *NTT1*-driven ATP import is directly mediated by functional mitochondria. In other words, high intracellular ATP levels become deleterious only when mitochondrial respiration is intact.

**Figure 7:**
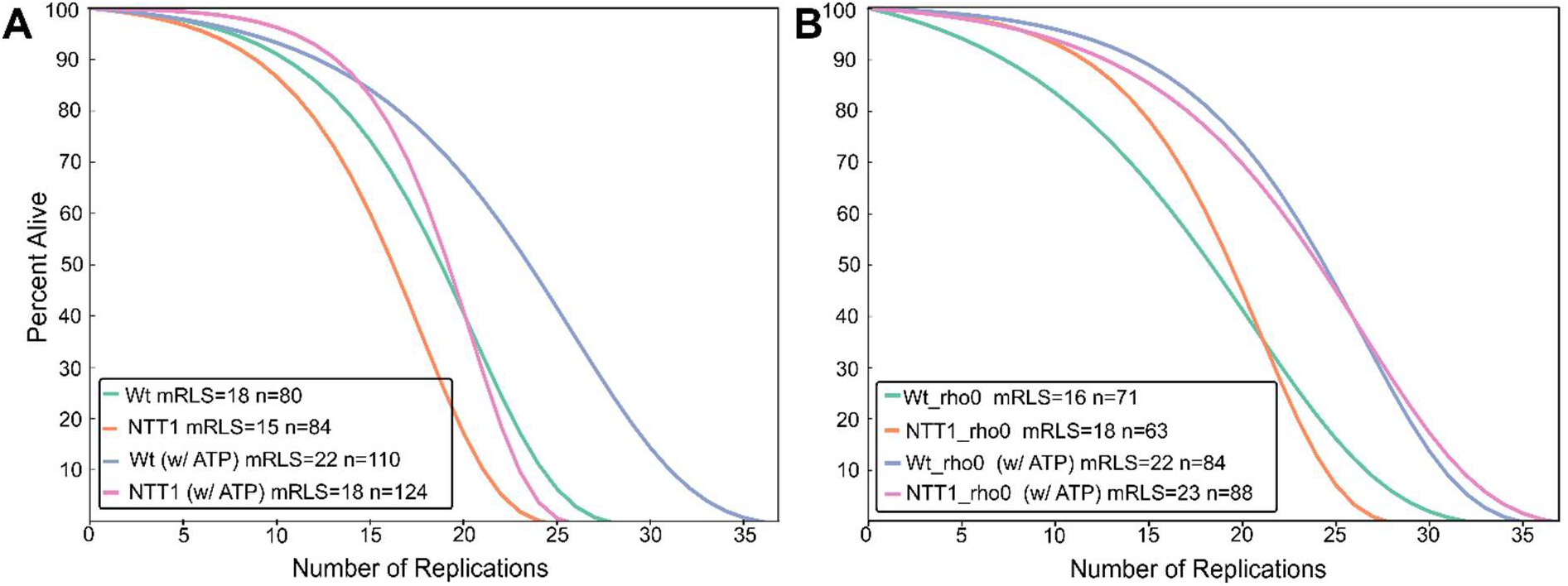
Replicative Lifespan analyses of *rho^0^* Isolates. Replicative lifespan (RLS) phenotypes of (**A**) parental controls and (**B**) their *rho^0^* isolates were measured using a microfluidic chip under untreated and 10 mM ATP-treated (w/ATP) conditions. Elimination of mtDNA in *rho^0^*isolates alleviates the toxic effect of ATP treatment on NTT1 cells. Figures indicate mean replicative lifespan (mRLS) values and number of individual mothers tested in each assay (n) were also included. Raw lifespan data and statistical significance are provided in **Supplementary File 3**.

Overall, these experiments allowed us to uncouple mitochondrial respiration from ATP-mediated lifespan effects. Strikingly, under baseline (no ATP treatment) conditions, the lifespan-shortening effect of *NTT1* expression was completely suppressed in *rho⁰* cells, suggesting that the toxicity of *NTT1*-driven ATP import requires functional mitochondria. In other words, high intracellular ATP becomes toxic only in the presence of intact mitochondrial function. Supporting this, at high extracellular ATP levels (10 mM), NTT1 cells showed a shortened lifespan, whereas *NTT1 rho⁰* isolates were protected and displayed significantly extended lifespan, again implicating mitochondria as mediators of ATP toxicity. Conversely, in wild-type cells lacking NTT1, extracellular ATP sensing (without import) robustly extended lifespan in both mtDNA-positive and *rho⁰* backgrounds, demonstrating that ATP sensing acts independently of mitochondrial function and does not confer toxicity.

These combined observations point to a dual mechanism: (i) extracellular ATP sensing promotes lifespan extension independently of mitochondrial respiration, and (ii) intracellular ATP import through *NTT1* affects lifespan in a mitochondria-dependent manner, with potential toxicity at high levels (**Fig. 8**). Thus, mitochondria serve as a key integrator of energy input via *NTT1*, while also distinguishing between beneficial signaling and detrimental metabolic overload. Additionally, the observation that ATP treatment extended lifespan in both Wt and Wt *rho^0^* isolates (**Fig. 7B**), indicating the positive lifespan effect of ATP is independent of mitochondrial function in Wt cells. In summary, the data supports a direct, synergistic relationship between *NTT1* function and altered ATP abundance in shaping mitochondrial function, with differential effects of extracellular ATP on Wt versus *NTT1* cells, where uncharacterized extracellular nucleotide sensing induces cellular changes that alter lifespan in yeast.

**Figure 8.**
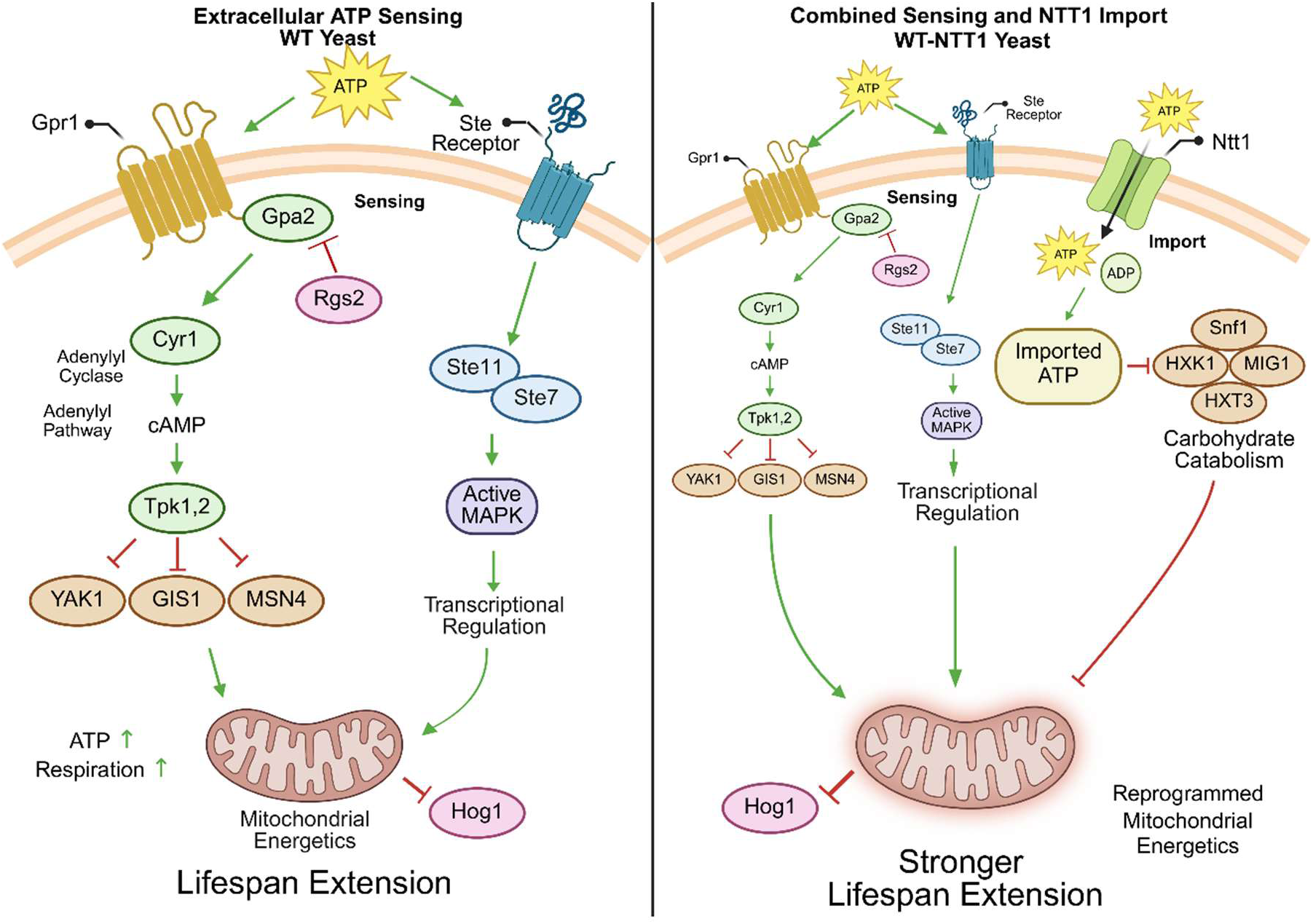
Proposed Models for the Effects of ATP Treatment in Wt and NTT1 Cells. The type- 1 effect involves extracellular ATP sensing, analogous to mammalian purinergic signaling, which activates G-protein signaling and regulates the stress-associated MAPK cascade to modulate mitochondrial bioenergetic function and downstream transcriptional responses (left panel). In contrast, the type-2 effect is specific to NTT1 cells and is driven by increased intracellular ATP abundance resulting from Ntt1-mediated ATP import from the medium. In this model, elevated intracellular ATP inhibits catabolic processes, reprograms mitochondrial bioenergetic function, and, in parallel, modulates some of the same pathways associated with the type-1 ATP-sensing response (right panel). In the type-2 effect, inhibition of Hog1 activity depends on the respiratory status of mitochondria. Both type-1 and type-2 effects increase replicative lifespan (RLS), with the type-2 effect producing a greater lifespan extension.

## Discussion

Catabolic systems (e.g., glycolysis, TCA cycle, respiration) are known to contribute significantly to cellular damage by generating by-products of cellular metabolism and reactive oxygen species (ROS) (54–56). A major purpose of catabolism is to generate ATP (55). However, downregulating subsets of these systems has been shown to extend lifespan in various organisms (57–60). Interestingly, as cells age, their ability to produce ATP often declines, leading to impaired cellular function and contributing to the aging process (61–63). The effect of altered ATP abundance in the context of aging remains largely unclear. This question has been difficult to address because ATP generation is essential for all life forms.

However, some eukaryotic intracellular parasites have evolved the ability to obtain ATP from their hosts. For example, *E. cuniculi* naturally uses nucleotide transporters to import ATP from the cytosol of eukaryotic host cells into its own cytosol as well as cellular compartments, such as mitochondria, to sustain its obligate intracellular lifestyle (28,29). We applied xenotopic synthetic biology strategy (64) to incorporate a nucleotide transporter from *E. cuniculi* into a budding yeast model and investigated the effects of altered ATP abundance on aging. Specifically, we expressed the *E. cuniculi* nucleotide transporter gene (*NTT1*) in yeast to enable ATP import from the extracellular environment, thereby aiming to downregulate much of the cellular catabolism while preserving energy supplies. This novel approach of directly providing energy to cells in the form of ATP alleviated organelle dysfunction and revealed a complex picture of metabolic and phenotypic alteration resulting in lifespan extension. Perhaps the most striking observation from our transcriptome data was that NTT1 cells downregulated glucose uptake and catabolic pathways, including energy generation and mitochondrial respiration, when supplemented with ATP. This observation suggests that when cellular energy demand is supported through a direct import of ATP, cells can adapt by suppressing carbohydrate catabolic processes to balance energy homeostasis. Furthermore, the observed concurrence of decreased expression of autophagy pathway genes and lifespan extension can be attributed to the fact that catabolic processes are a major source of cellular damage through the generation of reactive by-products (e.g., oxidants and secondary metabolites). Reduced catabolic activity may therefore generate fewer damaging agents, leading to decreased activation of damage-clearing pathways such as autophagy.

We found that overexpression of *NTT1* led to severe depletion of intracellular ATP and reduced lifespan. Follow-up experiments in cells lacking mitochondrial respiratory function revealed that this effect was linked to mitochondrial stress, as *rho^0^* isolates (lacking mitochondrial DNA and OXPHOS) of NTT1 cells restored lifespan to the level of Wt controls. In addition, while low-dose ATP treatment significantly increased intracellular ATP levels and extended lifespan, high-dose ATP supplementation was detrimental to lifespan in NTT1 cells but not in Wt cells. However, *NTT1*- *rho^0^* isolates did not display decreased lifespan under toxic ATP concentrations.

These observations strongly indicate that *NTT1* overexpression and altered ATP abundance interferes with mitochondrial function. Lifespan mode analysis under both treated and non-treated conditions further confirmed that age-associated decline in mitochondrial function can be compensated for by ATP supplementation, thereby promoting a positive lifespan effect. Our data showed that the percentage of cells aging through mitochondrial dysfunction (mode-2) was significantly reduced when the medium was supplemented with ATP. The majority of cells then followed a mode-1 aging trajectory, which is associated with a loss of Sir2 function. Interestingly, mode-1 cells also showed increased lifespan following ATP treatment. This suggests that elevated ATP levels are tightly connected to regulating cellular aging. In fact, previous studies demonstrated that overexpression of *HAP4* (a transcription factor regulating mitochondrial respiratory function) (65) alone did not extend lifespan in mode-2 cells. However, combined overexpression of *SIR2* and *HAP4* had a synergistic lifespan-extending effect, where cells lived significantly longer than those overexpressing *SIR2* alone (44). This supports the idea that mitochondrial function and Sir2 activity work together to modulate lifespan.

Remarkably, extracellular ATP sensing also exhibited life-extending effects following ATP treatment in Wt cells, but magnitude of the extension differed. Wt cells showed a median RLS increase of 41% (p = 1.28E-12) upon 10 µM ATP supplementation, while NTT1 cells showed an more pronounced extension (median RLS increase = 60%, p = 4.03E-18) in lifespan (Fig. 5D). Our transcriptomics data suggests the presence of a previously unknown nucleotide-sensing mechanism that links extracellular ATP to downstream cellular processes in yeast. Considering the altered expression and western blot analyses, we hypothesize that the Gpa2–cAMP–PKA pathway and stress responsive MAPK cascade may link extracellular ATP sensing to downstream pathways that promote mitochondrial respiration and increase ATP abundance. In fact, extracellular ADP treatment has also been shown to increase intracellular ATP levels in yeast (31). This suggests the existence of specific nucleotide-sensing mechanisms in yeast, similar to mammalian purinergic signaling, where purinergic receptors modulate downstream signaling cascades (66,67). A well-conserved *p38/HOG1* MAPK module in yeast is essential for cell survival under various stress conditions, such as osmotic, cell wall, nutrient, and oxidative stress (68–70). In addition, a wide variety of stimuli can elicit MAPK activation; however, it is still unclear how MAPKs regulate the distinct coupling of signals to downstream cellular responses (71,72). It is also known that MAPK is involved in cellular metabolic regulation and mitochondrial function and dynamics (73–75). Hence, our study suggests a previously uncharacterized MAPK–mitochondria signaling axis that regulates mitochondrial function in response to extracellular nucleotide sensing and provides new insights into the molecular and physiological adaptations of cells to ATP homeostasis. However, the components of the upstream sensing and downstream signaling cascades connecting ATP sensing to mitochondrial function require further investigation. It should also be highlighted that the increased mitochondrial respiration in these cells was not essential for the lifespan extension mediated by ATP supplementation, indicating modulation of other downstream canonical pathways. Our data also suggests that the presence or absence of respiratory active mitochondria might mediate alternative pathways that differentially regulate cellular lifespan following ATP treatment in NTT1 but not in Wt cells.

In part, this positive lifespan effect may also be associated with increased mitochondrial function and ATP abundance in Wt cells. However, we observed that Wt *rho^0^* isolates also showed significantly increased lifespan following ATP treatment, indicating that mitochondrial function is not essential for ATP signaling-mediated lifespan extension. It should be noted that caloric restriction (CR) also induces stress response pathways, increases respiration, and intracellular ATP abundance in yeast (76,77). However, CR-mediated lifespan extension is also independent of mitochondrial function, since Wt *rho^0^* isolates extend lifespan under CR conditions (78). This suggests that ATP-sensing mechanisms may regulate overlapping CR-mediated pathways to induce lifespan extension and may modulate alternative pathways independent of mitochondrial functional status. However, the exact mechanism of extracellular ATP-mediated lifespan extension and whether dual regulation of PKA and MAPK signaling through mitochondrial functional status are also required for CR mediated lifespan extension in yeast requires further exploration.

Overall, our study provides direct evidence that maintenance of cellular energy metabolism is a critical determinant of longevity and that preserving intracellular ATP levels can reduce entry into the mitochondrial dysfunction–associated aging trajectory (mode 2) in yeast. Declining mitochondrial function is widely recognized as a key hallmark of aging and has been supported by numerous previous studies. However, to our knowledge, this study is the first to directly demonstrate that modulation of ATP homeostasis itself can alter cellular aging. Our findings therefore establish cellular energy balance and mitochondrial function as tightly interconnected regulators of longevity and suggest that targeting mitochondrial bioenergetic capacity may represent a promising strategy for extending lifespan and preserving cellular health.

## Materials and Methods

### Yeast Strains and Plasmids

The Pleckstrin Homology (PH) domain of the *SLM1* gene was amplified from genomic DNA of the wild-type *S. cerevisiae* (BY4741) strain. Primers *PH-F* and *PH-R* were designed to amplify the PH domain with overlapping sequences compatible with assembly (primer sequences are listed in **Supplementary File 4**). The *NTT1* gene was amplified from a plasmid (28) using primers *NTT1-F* and *NTT1-R*, which included overlaps with the PH domain at the 5’ end and GFP at the 3’ end. The yeast-enhanced green fluorescent protein (yeGFP) sequence was amplified from the pYM25 plasmid (79) using primers GFP-F and GFP-R, incorporating overlaps with the *NTT1* sequence. The purified PH domain, *NTT1*, and yeGFP PCR products were fused using the NEBuilder HiFi DNA Assembly Master Mix (New England Biolabs, NEB #E2621).

The assembled *PH_NTT1_GFP* (**Supplementary File 4**) fusion construct was used as a template for PCR amplification with Gateway-compatible attB primers (*attB1-F* and *attB2-R*), which were designed to add recombination sites for Gateway cloning (primer sequences are provided in **Supplementary File 4**). The purified attB-flanked *PH_NTT1_GFP* fragment was recombined into the *pAG413GPD-ccdB* plasmid (Addgene plasmid #14142) (Alberti et al., 2007). The confirmed plasmid was transformed into BY4741 cells using the lithium acetate method.

A fusion cassette (**Supplementary File 4**) containing the PH sequence and *NTT1* (with a SmaI restriction site between them) was amplified by PCR from the appropriate DNA template using primers *Attb_PH_F* and *Attb_NTT1_R*. The reverse primer included a stop codon at the 3′ end of *NTT1*. The resulting attB-PCR product was purified and transferred into pAG306-GPD-ccdB (Addgene plasmid #41894). The final plasmid construct was integrated into the yeast genome after NotI digestion (Hughes et al., 2012). The empty pAG306-GPD plasmid (Addgene plasmid #41895) was integrated into the yeast genome as a control strain.

Finally, the yeast strains containing the QUEEN ATP reporter gene were obtained from the National BioResource Project Yeast Genetic Resource Center. The *PH_NTT1* construct (with a stop codon) in pAG306-GPD-ccdB was integrated into the BY29034 strain. The empty pAG306- GPD plasmid was also integrated into the BY29034 genome as a control strain. All sequences were verified by Sanger sequencing after PCR amplification and cloning.

### RNA Isolation and RNA-seq Analysis

Cells (three independent cultures for each strain) were cultivated in Complete Synthetic Medium (CSM) at 30°C with constant agitation (200–220 rpm) until they reached an optical density at 600 nm (OD600) of approximately 0.4–0.5. To examine the effect of extracellular ATP, a 50 mM ATP stock solution was prepared in CSM to maintain a consistent medium composition during treatment. Cells were treated with ATP at a final concentration of 5 mM for 1 hour at 30°C under continuous shaking. Untreated control cultures of the same strains were grown under identical conditions but without ATP supplementation.

After treatment, cells were harvested by centrifugation at 4°C. Total RNA was extracted using the Quick-RNA Fungal/Bacterial MiniPrep™ Kit (Zymo Research R2014) according to the manufacturer’s instructions. The resulting RNA samples were used for downstream RNA-sequencing analysis.

To prepare RNA-seq libraries, the Illumina TruSeq RNA library preparation kit was used according to the user manual. RNA-seq libraries were loaded onto the Illumina NovaSeq 6000 platform to produce 150 bp paired-end sequences. After quality control and adapter removal, paired-end clean reads were aligned to the yeast reference genome (Ensembl, R64-1-1) using Hisat2 v2.0.5. FeatureCounts v1.5.0-p3 was used to count the number of reads mapped to each gene. Fragments per kilobase of transcript per million mapped reads (FPKM) values were calculated based on the length of the gene and the read count mapped to that gene.

Differential expression analysis was performed using the DESeq2 R package (v1.20.0). The resulting p-values were adjusted using the Benjamini and Hochberg approach for controlling the false discovery rate. Genes with an adjusted p-value ≤ 0.05 were classified as differentially expressed. Gene Ontology (GO) enrichment analysis of differentially expressed genes was implemented using the clusterProfiler R package, which corrects gene length bias. GO terms with corrected p-values less than 0.05 were considered significantly enriched among differentially expressed genes.

### Microscopy for NTT1 Localization

The *S. cerevisiae* strain harboring *NTT1* was grown in yeast peptone dextrose (YPD) medium at 30°C with constant agitation (200–220 rpm) until mid-log phase. For visualization of *NTT1* localization, cells were harvested and immediately examined by confocal fluorescence microscopy using a GFP-compatible excitation/emission range (e.g., 488 nm excitation/505–530 nm emission). Images were acquired under identical settings for all samples to enable direct comparisons of fluorescence signals.

### Relative ATP Measurement Using QUEEN ATP Reporter and LC-MS/MS

The *S. cerevisiae* strains expressing the QUEEN ATP reporter (NTT1 and Wt control) were cultivated in Complete Synthetic Medium (CSM) at 30°C with constant agitation (200–220 rpm) until reaching mid-log phase (OD600 ∼ 0.4–0.6). The cells were retained in CSM for one set of experiments and either treated with 5 mM ATP for 30 minutes or left untreated as a control.

In a separate experiment, cells were transferred to CSM supplemented with 20 mM 2DG (Sigma #D-8375) for 30 minutes to modulate intracellular ATP levels. Following the indicated treatments, cells were harvested, washed once in fresh CSM, and resuspended in CSM for microscopy. Cell viability was assessed using propidium iodide (PI) staining.

Images were acquired using a Leica DMi8 inverted fluorescence microscope equipped with 405, 488 and 561 nm lasers, an LED3 fluorescence illumination system, a 100X, 1.47 numerical aperture Plan-Apochromat oil immersion lens, a Flash 4.0 v3 sCMOS camera (Hamamatsu), a W-View Gemini beam splitting device (Hamamatsu), and LAS X v3.7.6.25997 software. All strains within an experiment were imaged under identical exposure settings. A series of four images were captured for each field: one bright-field image and three fluorescence images corresponding to excitation/emission filters optimized for QUEEN (QWF-T ex405, QWF-T ex488), or for PI (QWF-T ex561).

Digital images were processed using Fiji software. First, the background signal was subtracted from the fluorescence channels after converting images to 32-bit floating-point format. Intracellular ATP was estimated by calculating a QUEEN ratio, defined as the pixel intensity of the ex405 image divided by the corresponding intensity of the ex488 image on a per-pixel basis. The mean QUEEN ratio for each cell was determined by averaging pixel ratios within a manually defined region of interest (ROI) encompassing the cell. Higher QUEEN ratios indicate elevated intracellular ATP levels. PI fluorescence (captured via ex561) was used to identify and exclude nonviable or compromised cells from the analysis. Based on the normality and homogeneity assumptions (Shapiro-Wilk and Levene’s tests, respectively), a non-parametric Kruskal-Wallis test was chosen for comparison of groups. Post hoc pairwise comparisons using the Wilcoxon rank sum test with Bonferroni adjustment were used when comparing only two groups.

For the LC-MS/MS analyses, yeast cultures of both the NTT1 cells and the Wt strain were grown in liquid medium at 30 °C with shaking at 210 rpm until mid-logarithmic phase (OD₆₀₀ ≈ 0.4). A 50 mM ATP stock solution was freshly prepared from ATP disodium salt (Cayman Chemical, Cat. No. 21121). Cultures were divided into experimental groups consisting of untreated controls, 1 mM ATP, and 5 mM ATP supplementation. ATP was added to cultures at the indicated concentrations, and cells were incubated for 30 min before harvesting. Cells were collected by centrifugation at 5000 rpm for 10 min, washed three times in PBS, and quenched in 500 µL of 80% methanol (–20 °C). Quenched cells were homogenized by tip sonication for 30 seconds in an ice bath, followed by a 30-second pause, and the cycle was repeated three times. Lysates were transferred to 4 mL glass vials, and the tubes were rinsed with an additional 200 µL of 80% methanol, which was combined with the extract. Subsequently, 1.4 mL of cold chloroform (–20 °C) was added, and the mixture was vortexed for 30 seconds and shaken at 300 rpm for 10 min at 4 °C. After the addition of 700 µL of cold water, samples were shaken vigorously for 1 min and centrifuged at 3000 rpm for 5 min at 4 °C. The upper aqueous phase (1 mL) was collected, dried using a SpeedVac concentrator for approximately 3 h, and stored at –80 °C. Prior to LC-MS/MS measurement, dried extracts were reconstituted in 100 µL of formic acid in water containing isotopically labeled internal standards (Metabolomics QReSS, Cambridge Isotope Laboratories). A pooled quality control (QC) sample was prepared by combining 15 µL from each extract, and 10 µL was injected per run. Orbitrap IDX Tribrid mass spectrometer (Thermo Fisher Scientific) coupled with a Vanquish UHPLC system was used for detecting the ATP levels. Separation was achieved on a Waters BEH C18 column (2.1 × 150 mm, 1.8 µm) maintained at 30 °C with a flow rate of 250 µL/min. Raw LC-MS/MS data were processed in MS-DIAL v4.92 (Tsugawa et al., 2015). Blank samples were included to identify and remove background signals. Similar methodology was applied to measure ADP and AMP levels from untreated cells. Each strain was processed and analyzed independently, and all subsequent metabolite extraction, LC-MS/MS acquisition, and data processing were carried out separately on different days.

### Immunoblotting Bioassays

Hog1 phosphorylation was analyzed using antibodies that recognize the total and phosphorylated forms. Mid-log phase cells were grown on YPD medium, treated or not treated with 5mM ATP. Proteins were extracted using yeast protein extraction buffer YPER (Thermo, Catalog number 78990) supplemented with protease and phosphatase inhibitors mix (Thermo, Catalog number 87785). Whole cell extracts were resuspended in boiling SDS-PAGE sample buffer for 5 min. Two independent biological replicate experiments were performed using distinct phospho-p38 antibodies to confirm the result. In the first experiment, membranes were probed with phospho- p38 (Hog1) antibody at 1:1000 (Cell Signaling Technology, 9215T), Hog1 at 1:1000 (Santa Cruz, sc-165978), and Pgk1 at 1:5,000 (Abcam, 113687). In the second, independent experiment, a full new set of biological replicates was grown, treated, and processed as described above, and membranes were probed with a different phospho-p38 antibody. Phospho-p38 MAPK (Thr180/Tyr182) polyclonal antibody at 1:1000 (Proteintech, 28796-1-AP), along with total Hog1 and Pgk1 antibodies as mentioned above. For both experiments, following incubation with secondary antibodies, membranes were developed using chemiluminescent substrate (Bio-Rad, 1705060) and visualized with a Bio-Rad imaging system. Band intensities were quantified in Fiji/ImageJ (v.2.14.0) from 16-bit captures with no brightness, contrast or levels adjustment applied prior to analysis. For each lane, a rectangular region of interest of fixed dimensions was positioned over the band, and local background was estimated from band-free regions of identical width immediately above and below the band within the same lane and subtracted as a linear ramp. Band intensity is reported as background-subtracted integrated density in arbitrary units. Phospho- Hog1 was normalised to total Hog1 in the same lane, and total Hog1 to Pgk1; all values are expressed relative to untreated wild-type cells. All lanes within a panel derive from a single membrane imaged in a single exposure; no lanes were cropped or spliced.

### Setting up Microfluidics Experiments

The microfluidics device fabrication and experimental setup have been described previously (46,48).Yeast cells were inoculated into 2ml synthetic complete medium (SD, 2% glucose) and incubated on a 30 ^0^C shaker overnight. From this culture, 2 µL of saturated culture was transferred to 20ml SD and incubated until its OD600 reached ∼0.6 (early exponential growth phase). A microfluidics chip containing 4 microfluidics channels was placed under vacuum for at least 15min. In the meanwhile, 50ml SD with 0.04% Tween-20 (Sigma) was transferred to a 60ml syringe, to which plastic tubing (TYGON, ID 0.020 IN, OD0.060 IN, wall 0.020 IN) was connected. For samples that have additional ATP in the medium, ATP was added to this 50ml medium to reach the corresponding concentrations. The syringe was sealed with aluminum foil. 4 such syringes were prepared, 1 for each microfluidics channel. After vacuum, the microfluidics chip was quickly connected to the syringes through plastic tubing from its inlet port to avoid air flowing into the microfluidics channels. The outlets were then connected to plastic tubing filled with ddH2O. The chip with inlets and outlets was then fixed to a microscope stage. For cell loading, cells were transferred into a 60ml syringe with plastic tubing connected. Connection of the microfluidics inlets to the media supply syringe was then replaced by connection to the cell loading syringe. The flow of medium in the device was maintained by gravity to drive cell loading into the traps. After loading, the media supply tubing was switched back to the inlet. Heights of all supply syringes were then adjusted to have a height difference between inlet and outlet of around 240cm. Tween- 20 is a non-ionic surfactant that helps reduce friction during cell loading. We have validated previously that this low concentration of Tween-20 has no significant effect on cellular lifespan or physiology (45).

As a circular permutated GFP, the QUEEN reporter can be excited by both 410nm and 488nm excitation wavelengths. When bound by ATP, the 410nm excitation yields higher fluorescence, while the 488nm excitation yields lower fluorescence, and vice versa without ATP (30, 31). Therefore, intracellular ATP levels are typically quantified using the ratio of fluorescence intensities obtained from these two excitation wavelengths (410nm/488nm). In our aging experiments, repeated and prolonged exposure to the high-intensity 410nm excitation causes phototoxicity in yeast cells and impacts yeast lifespan. We thus only measured the QUEEN reporter fluorescence at the 488nm excitation for aging experiments and plotted the inverse of fluorescence measurements (1/488nm), as an indicator of ATP abundance, in Fig. 5, A-C.

For assaying rho0 isolates, yeast cells were cultured overnight in synthetic complete (SC) media supplemented with 2% glucose. The following day, cultures were diluted 1:300 into 4 mL fresh SC media and incubated at 30°C for 3 hours. Prior to loading into the microfluidic device, 0.05% bovine serum albumin (BSA) was added to the culture to reduce nonspecific adhesion. Cells were then introduced into the microfluidic device, and all experiments were conducted using a Nikon Eclipse Ti2 inverted microscope equipped with a 60× 1.4 NA oil immersion objective (Nikon) and maintained at 30°C using a stage-top incubation system (Okolabs). Media was continuously delivered to the device via two 30 mL syringe pumps (total flow rate of 4 µL/min) to ensure nutrient availability and waste removal. Image acquisition was carried out every 5 minutes for up to 72 hours using NIS-Elements software, with the Nikon Perfect Focus System (PFS) engaged throughout all imaging sessions to preserve focal stability. Cell viability and death events were monitored in real time, and all downstream image segmentation, cell tracking, and data extraction were performed using MATLAB (MathWorks).

### Time-lapse Microscopy

Time-lapse Microscopy experiments were conducted using a Nikon Ti-E inverted fluorescence microscope with an EMCCD camera (Andor iXon X3 DU897). The light source is a spectra X LED system. Images were taken using a CFI plan Apochromat Lambda DM 60X oil immersion objective (NA 1.4 WD 0.13mm). In the yeast aging experiments, images were taken for each imaging channel every 15min for a total of 90 hours. The setting of each channel was set as the following and remains consistent across time-lapse imaging aging experiments: Phase, 60ms; GFP, 100ms at 10% lamp intensity with an EM gain of 100.

### Quantification of Single-cell Aging Traces

The procedure of image quantification has been described previously (48). Image processing was done using ImageJ and custom Matlab codes. The background of fluorescence channel was subtracted by rolling ball background subtraction. Cells were masked and tracked by thresholding the GFP channel. The correctness of cell tracking was manually checked. The mean intensity of the top 40% pixels was quantified for each mother cell at each time point. Because the QUEEN reporter was not uniformly distributed in aged yeast cells, whose enlarged vacuoles lack the reporter, we selectively analyzed the brightest pixels to exclude low-intensity regions unrelated to ATP levels. Cell aging mode was determined by daughter cell morphology (44). Cell replicative lifespan was manually quantified. Cells that showed obvious abnormal morphology upon loading were filtered out of RLS analysis.

## Supporting information

SIle S1

## Acknowledgments

We thank Dr. Robert P. Hirt (Newcastle University) for providing plasmids containing the cDNA-based expression cassette of *NTT1*, and Dr. Satoshi Yoshida (Waseda University, Japan) for providing strains and plasmids harboring the QUEEN ATP reporter vector through the National BioResource Project (NBRP-Japan) Yeast Genetic Resource Center. We also thank the NBRP staff for handling the collection and preparing and sending these strains. Additionally, we acknowledge the support provided by The Cancer Center Shared Resource for Metabolomics & Lipidomics at the University of Virginia for performing LC-MS/MS analyses.

## Data Availability

All data generated or analyzed during this study are included in the manuscript and supporting files. RNA-seq data were deposited with the NCBI Gene Expression Omnibus (GEO) with accession number GSE312907.

## Author contributions

A.K. and V.N.G. designed research; N.O., H.S., V.S., P.P., R.H., J.H.S., D.C.P., performed research.

N.O., H.S., V.S., P.P., R.H., J.H.S., V.M.L., N.H., and A.K. analyzed data; and N.O., H.S., V.S., P.P., V.M.L., D.C.P., V.N.G., N.H., and A.K, wrote the paper.

---

**Supplementary Figure 1.**
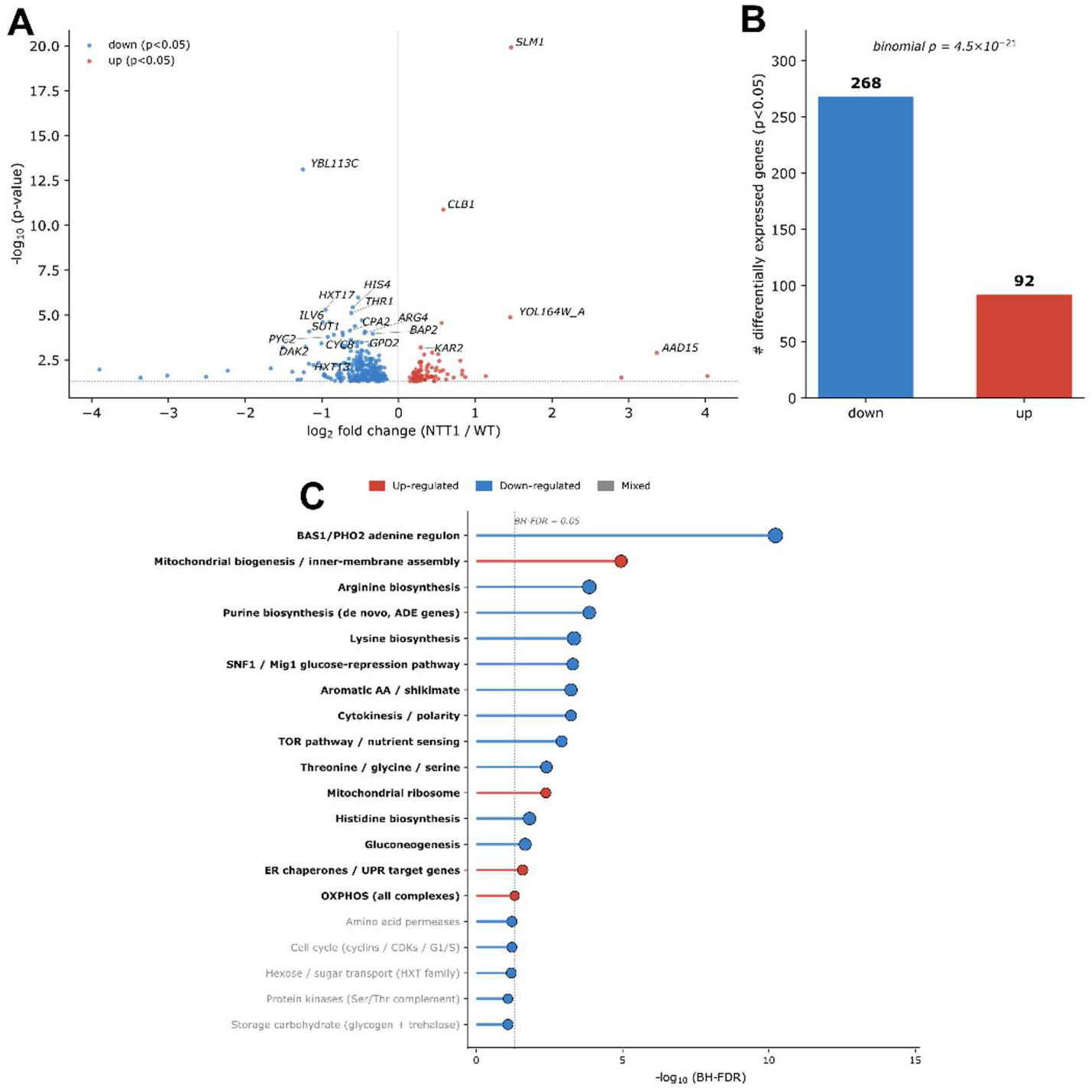
Differential expression in NTT1-overexpressing cells. (**A**) Volcano plot of log2 fold change vs −log10 nominal p-value for all expressed genes. Blue and red points denote genes passing p < 0.05 with negative or positive fold changes, respectively. Genes with the largest effect sizes or strongest statistical support are labelled. (**B**) Numbers of significantly down- and up-regulated genes (p < 0.05); the asymmetry (268 down vs 92 up) is significant by binomial test against an unbiased null. (**C**) Functional enrichment of NTT1 vs Wt differential expression. Categories with BH-FDR < 0.05 are shown, ranked by significance. Red, predominantly up- regulated; blue, predominantly down-regulated; circle size scales with fold enrichment. Numbers in parentheses are differentially expressed genes / total in category. Curated gene sets cover central metabolic, signaling and biogenesis pathways in S. cerevisiae.

**Supplementary Figure 2.**
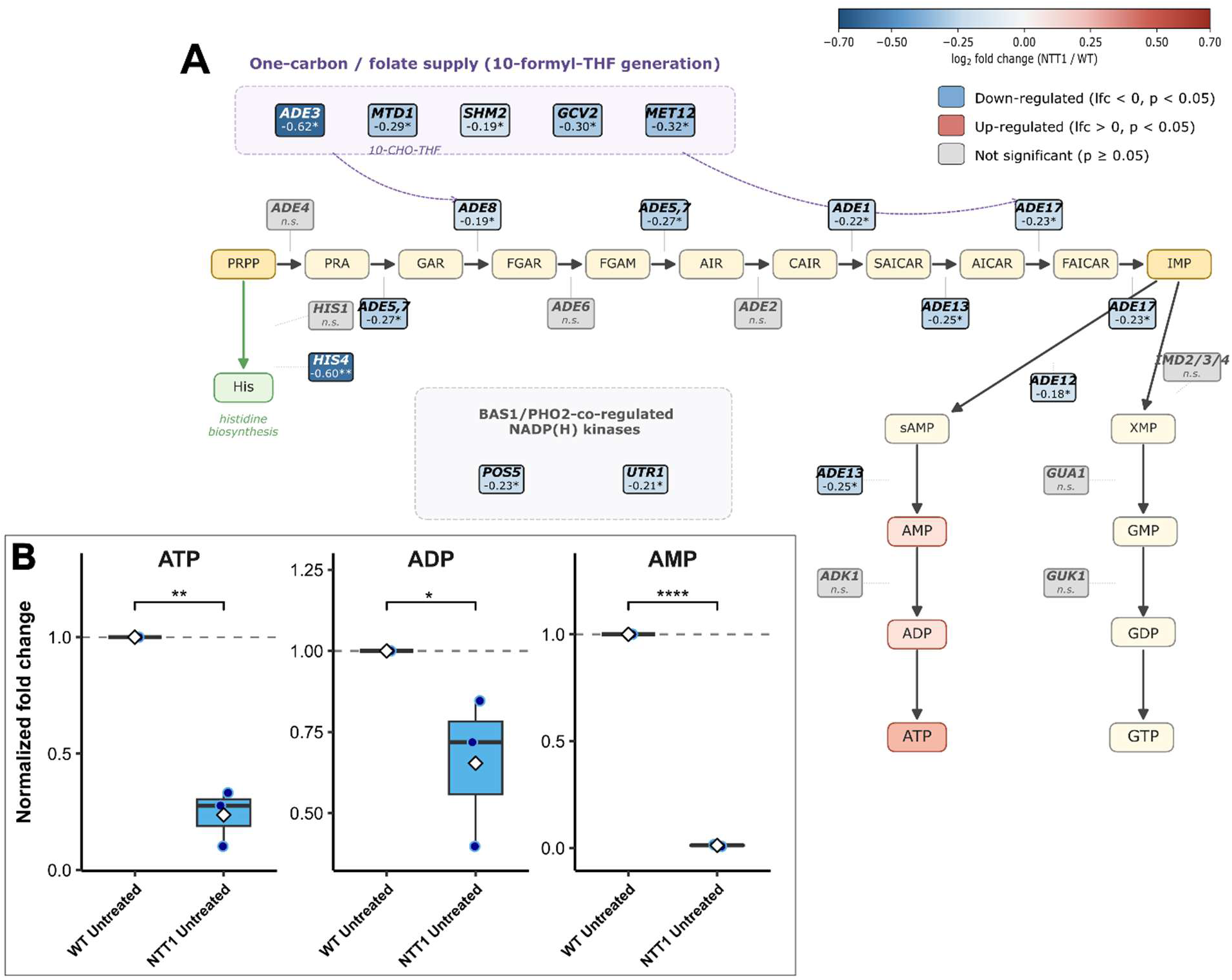
Purine specificity of the NTT1 response. (**A**) Log2 fold changes of de novo purine (ADE), BAS1-co-regulated (HIS4) and one-carbon/folate (MTD1, SHM2, GCV2, ADE3) genes that reached p < 0.05. Asterisks indicate p < 0.05. All thirteen captured genes are down-regulated. None of the de novo pyrimidine (URA) or ribonucleotide reductase (RNR) genes reached p < 0.05; the parallel pathways are transcriptionally unaffected. (**B**) The LC-MS/MS measurement of purine metabolites in Wt versus NTT1 cells. The raw LC-MS/MS data along with statistical values are available in **Supplementary File 1**.

**Supplementary Figure 3.**
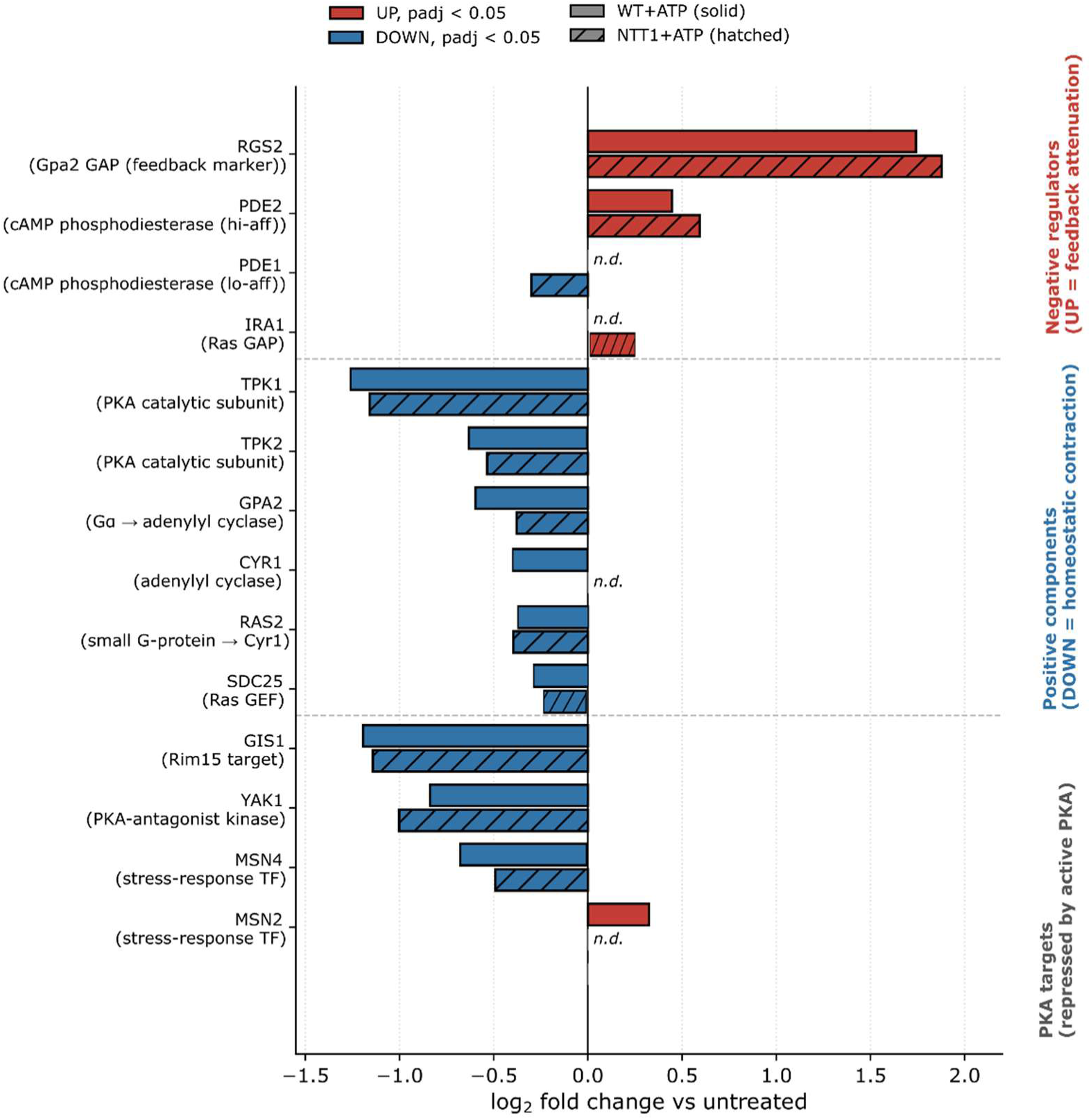
Extracellular ATP treatment activates Gpa2–cAMP–PKA pathway signaling. Log2 fold change (versus untreated) for Gpa2–cAMP–PKA pathway genes in Wt (solid) and NTT1 cells (hatched) after ATP treatment. Red, upregulated; blue, downregulated (padj < 0.05); n.d., not significant. Upregulation of negative regulators alongside repression of positive components and PKA targets indicates active PKA signaling in both strains.

**Supplementary Figure 4:**
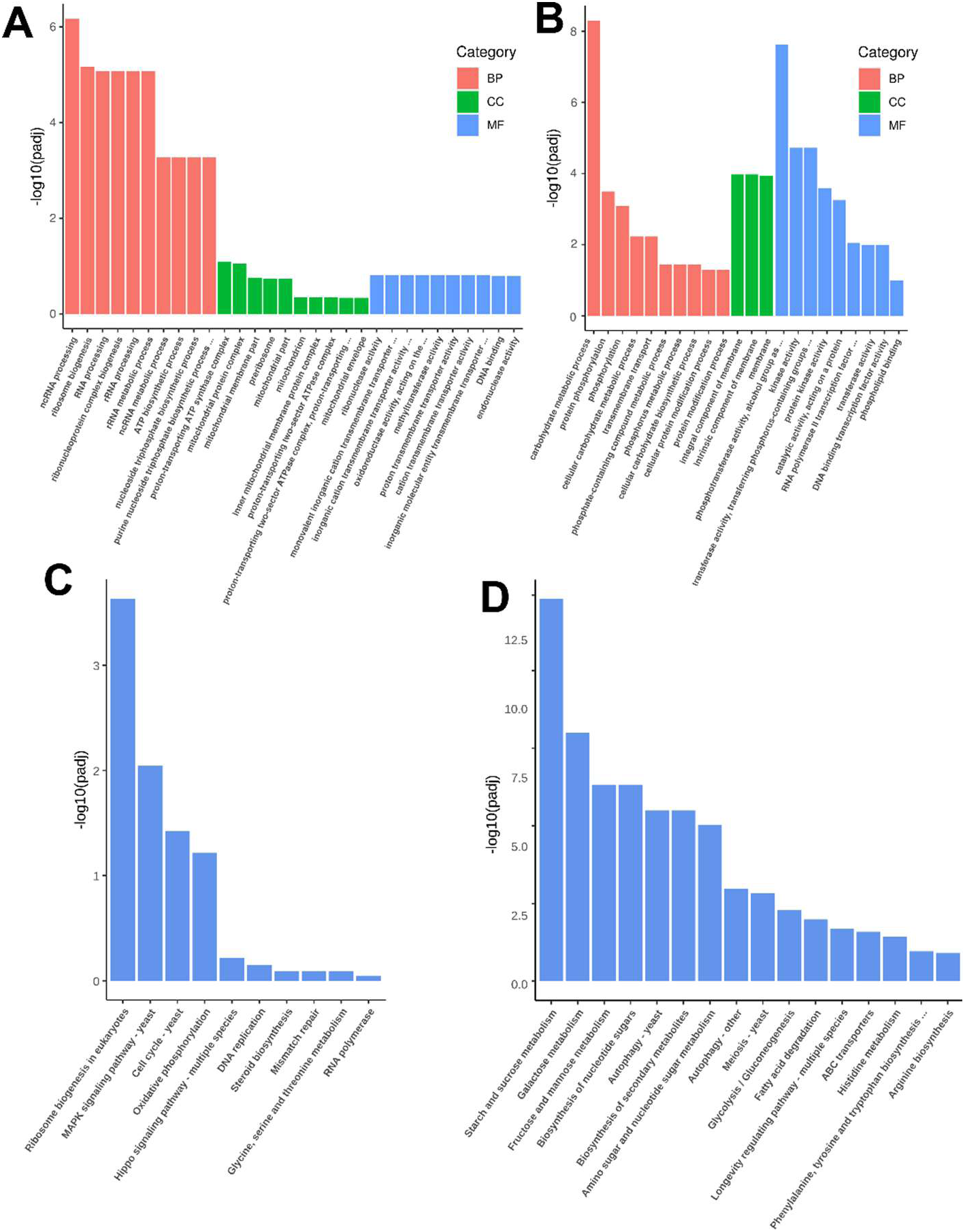
Gene Ontology and KEGG Pathway Analyses for Wt Cells. Gene ontology (GO) enrichment analysis for (**A**) upregulated and (**B**) downregulated genes, and KEGG pathway analysis for (**C**) upregulated and (**D**) downregulated genes in Wt cells after ATP treatment. GO categories include BP (Biological Process), CC (Cellular Component), and MF (Molecular Function).

**Supplementary Figure 5:**
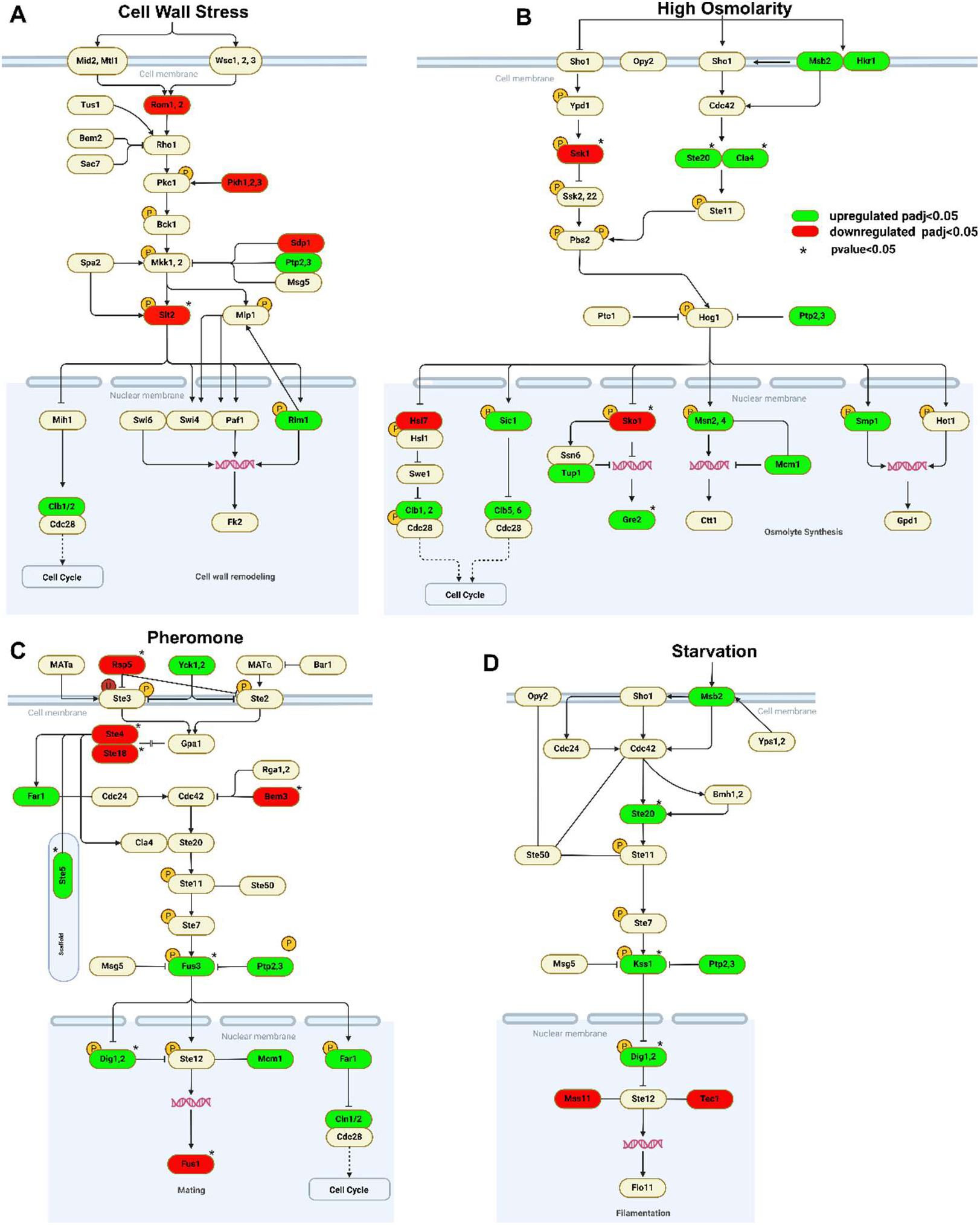
Alteration of MAP Kinase Pathway in Wt Cells. Genes uniquely upregulated (green) or downregulated (red) are shown for each component of the MAPK pathway in Wt cells under ATP treatment: (**A**) Cell Wall Stress, (**B**) High Osmolarity, (**C**) Pheromone, and (**D**) Starvation.

**Supplementary Figure 6.**
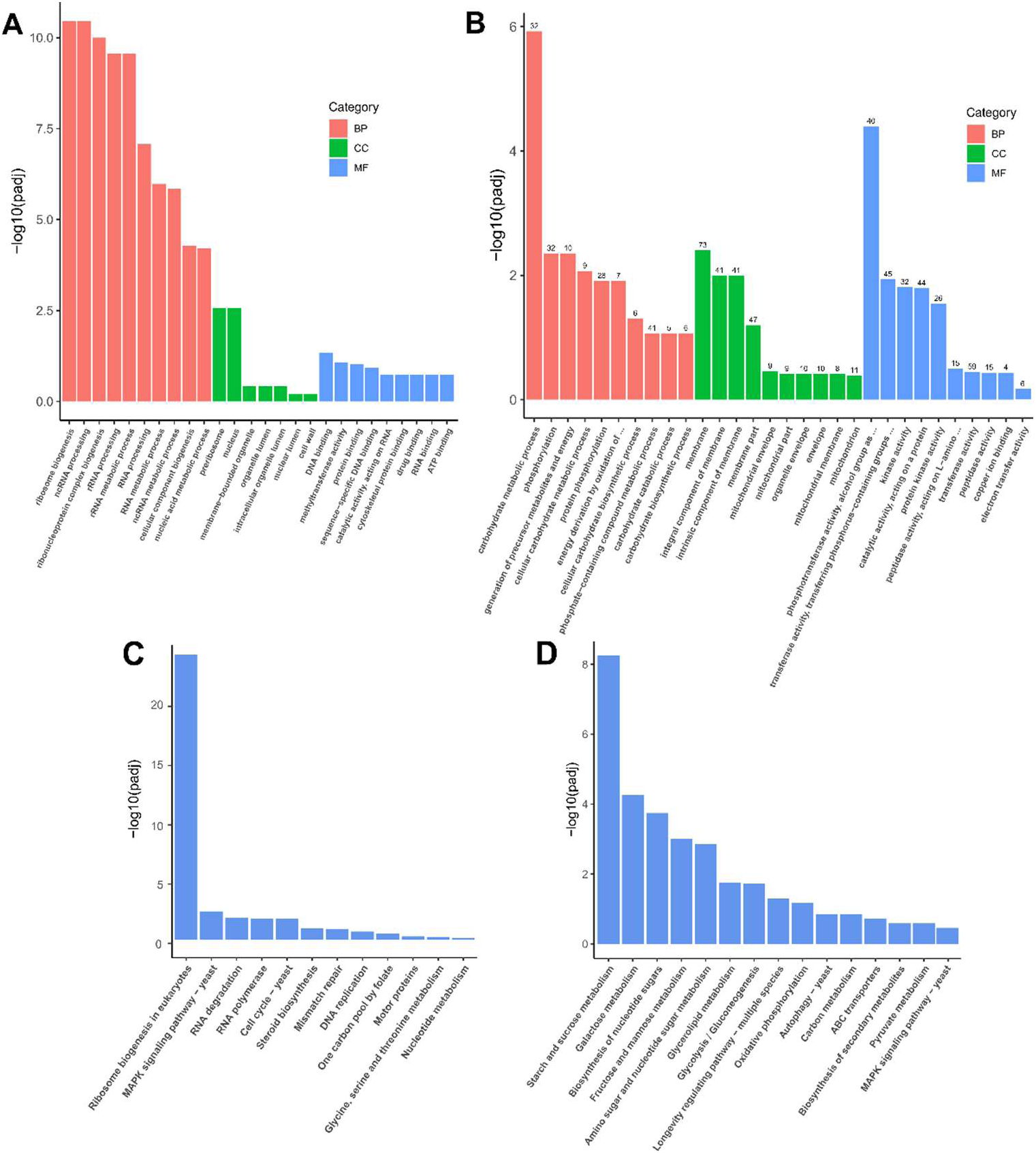
Gene Ontology and KEGG Pathway Analyses for NTT1 Cells. Gene ontology (GO) enrichment analysis for (**A**) upregulated and (**B**) downregulated genes, and KEGG pathway analysis for (**C**) upregulated and (**D**) downregulated genes in NTT1 cells after ATP treatment compared to untreated controls. GO categories include BP (Biological Process), CC (Cellular Component), and MF (Molecular Function).

**Supplementary Figure 7.**
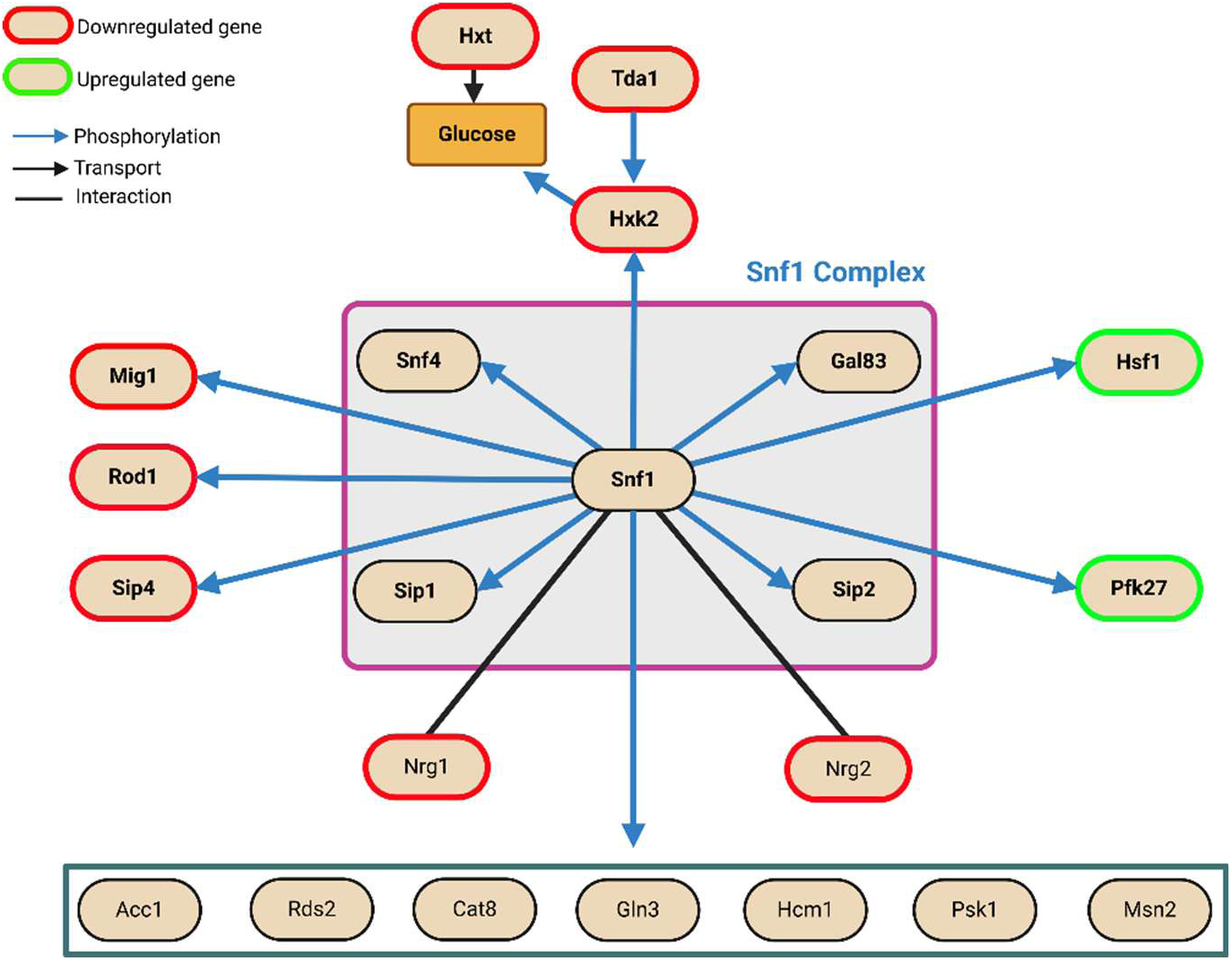
Snf1-Mediated Regulatory Gene Network in NTT1 Cells. Genes regulated by Snf1 in NTT1 cells under ATP treatment are depicted in red (downregulated), green (upregulated), and black (no expression changes). The types of interactions are represented by arrows and lines.

**Figure S8:**
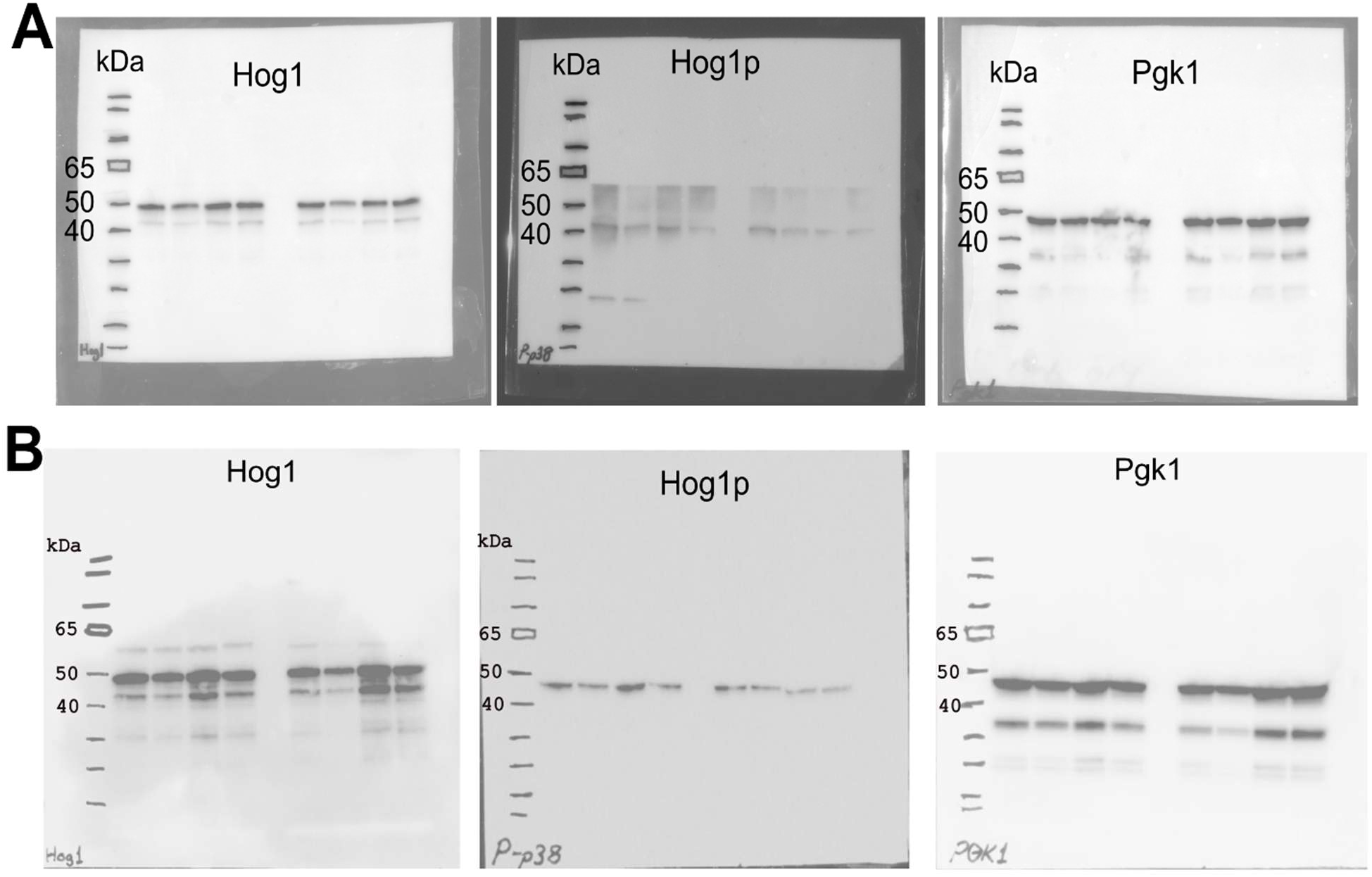
Uncut western blot membranes for assessing the active abundance of Hog1 protein with two independent antibodies raised against phospho-Hog1 (Hog1^P^).

## Supplementary Files

**Supplementary File 1:** Raw data for ATP, ADP, and AMP measurements obtained using the QUEEN reporter and LC-MS/MS.

**Supplementary File 2**: Raw RNA-seq counts and FPKM values, along with log2 fold-change values.

**Supplementary File 3:** Raw replicative lifespan measurements and single-cell ATP levels for Wt and NTT1 strains.

**Supplementary File 4:** NTT1 coding sequence.

