## Supplementary material for "Engineering ATP Import in Yeast Uncovers a Synthetic Route to Extend Cellular Lifespan": File S4

**PH_NTT1**

**atg**AGGACATTCAAAGAAATCGTTTACAAATACCAATTCGATCCTTTGACTTATGAAATCAAATCAGGGTTTTTAGAAAGAAGATCAAAATTTCTAAAATCCTATTCAAAAGGGTATTATGTGCTGACACCGAACTTTTTACATGAATTCAAGACCGCGGACAGAAAAAAAGATTTAGTACCAGTGATGTCGTTAGCATTAAGTGAATGTACTGTGACGGAACATTCCCGGAAAAACAGTACATCATCACCAAATTCGACTGGTTCGGATGCTAAGTTTGTGTTGCATGCTAAGCAAAACGGCATAATTCGTCGTGGTCACAATTGGGTTTTTAAAGCCGATTCTTATGAATCAATGATGTCTTGGTTTGATAATTTGAAGATCTTAACATCTACTTCGAATATACAAGACAAG**CCCGGG**AATGAGGTGGAAAACAACAACCACAGTTTTCCTCGTGAAGACATCCCCACAGAAGACGAGATTGAAGAAGAGGCCAACAGCAGGCAGGGAATCCTCCGCTACTTCAGGGTGGCAAGAGCAGAATACACCAAGTTCGCTCTCTTGGGCCTGATGTTTGGAATAATCGGATTTATATACTCGTTCATGAGAATTCTCAAGGACATGTTTGTGATGGTCAGGCAGGAGCCCACGACAATATTGTTCATCAAGATCTTCTACATCCTTCCAGTCTCAATGGCCCTGGTCTTCCTCATACAGTACATGCTGGGGACAAAAACAGTCTCAAGGATATTCTCCATCTTTTGTGGAGGATTTGCATCCCTATTCTTTCTGTGTGGCGCAGTGTTTCTGATAGAGGAGCAGGTCTCTCCCTCGAAGTTCCTGTTCAGAGACATGTTTATAGACGGAAAAATGTCCAGCAGAAGCCTCAATGTCTTCAAGTCGATGTTTCTGACACTCAACGAGCCCCTCGCAACAATCGTCTTCATTTCAGCCGAGATGTGGGGAAGCCTTGTTCTGTCGTATCTCTTCCTCAGCTTCCTGAACGAGTCATGCACGATTAGACAGTTCTCTAGGTTCATTCCTCCCCTCATAATCATCACAAACGTGTCGCTTTTCCTCTCGGCAACGGTGGCAGGAGCCTTTTTCAAGCTCCGGGAGAAGTTAGCATTCCAGCAGAATCAGGTCCTGCTCTCGGGAATCTTCATCTTCCAGGGCTTTCTGGTGGTCCTGGTGATCTTCTTAAAGATATACCTCGAAAGAGTGACAATGAAGAGGCCCCTGTTCATCGTCTCCTCGGGGTCAAGGAGAAAGAAGGCCAAGGCCAACGTGTCCTTTTCAGAAGGCCTTGAGATCATGTCCCAGTCAAAGCTGCTGCTTGCAATGTCCCTCATCGTCCTCTTCTTCAACATCTCCTACAACATGGTCGAGTCCACATTCAAAGTCGGAGTTAAGGTTGCTGCAGAATACTTCAACGAGGAAAAGGGGAAATACTCCGGAAAGTTCAACCGGATCGACCAGTATATGACCTCTGTCGTGGTCATATGCCTCAACCTGTCACCATTCTCGAGCTATGTCGAGACAAGAGGGTTTCTTCTGGTCGGGCTTATAACGCCTATTGTGACACTCATGGCCATCGTTCTTTTCCTAGGGTCTGCGCTTTACAACACTAGCATGGAAGAGTCTGGGCTTGGAATCGTGAACGGGCTGTTTCCAGGTGGAAAGCCCCTGTACGTCCTCGAGAACTACTTCGGCGTAATCTTCATGTCGCTTCTGAAGATCACGAAGTACTCTGCATTTGACATCTGCAAAGAAAAGCTTGGGATGCGGATCAACCCCACATACAGAGCAAGGTTCAAGAGCGTGTATGATGGGATTTTTGGGAAGCTTGGGAAGTCCATAGGATCGATATACGGGCTCCTGATGTTCGAGGCCTTGGACACGGAAGACCTTAGAAAGGCAACGCCGATAACAGCAGGGATTATCTTTATCTTCATTGTCATGTGGGTAAAAGCGATAATCTACCTGTCGAGGTCTTACGAGTCTGCTGTACAGCATAATAGAGATGTTGACATCGACATGACCGAAAAGGCAAAGAAAAGCTTGGAGACTCCTGAGGAGCCAAAAGTTGTAGAT**taa**

**PHNTT1GFP**

*ATGAGGACATTCAAAGAAATCGTTTACAAATACCAATTCGATCCTTTGACTTATGAAATCAAATCAGGGTTTTTAGAAAGAAGATCAAAATTTCTAAAATCCTATTCAAAAGGGTATTATGTGCTGACACCGAACTTTTTACATGAATTCAAGACCGCGGACAGAAAAAAAGATTTAGTACCAGTGATGTCGTTAGCATTAAGTGAATGTACTGTGACGGAACATTCCCGGAAAAACAGTACATCATCACCAAATTCGACTGGTTCGGATGCTAAGTTTGTGTTGCATGCTAAGCAAAACGGCATAATTCGTCGTGGTCACAATTGGGTTTTTAAAGCCGATTCTTATGAATCAATGATGTCTTGGTTTGATAATTTGAAGATCTTAACATCTACTTCGAATATACAAGACAAG*AATGAGGTGGAAAACAACAACCACAGTTTTCCTCGTGAAGACATCCCCACAGAAGACGAGATTGAAGAAGAGGCCAACAGCAGGCAGGGAATCCTCCGCTACTTCAGGGTGGCAAGAGCAGAATACACCAAGTTCGCTCTCTTGGGCCTGATGTTTGGAATAATCGGATTTATATACTCGTTCATGAGAATTCTCAAGGACATGTTTGTGATGGTCAGGCAGGAGCCCACGACAATATTGTTCATCAAGATCTTCTACATCCTTCCAGTCTCAATGGCCCTGGTCTTCCTCATACAGTACATGCTGGGGACAAAAACAGTCTCAAGGATATTCTCCATCTTTTGTGGAGGATTTGCATCCCTATTCTTTCTGTGTGGCGCAGTGTTTCTGATAGAGGAGCAGGTCTCTCCCTCGAAGTTCCTGTTCAGAGACATGTTTATAGACGGAAAAATGTCCAGCAGAAGCCTCAATGTCTTCAAGTCGATGTTTCTGACACTCAACGAGCCCCTCGCAACAATCGTCTTCATTTCAGCCGAGATGTGGGGAAGCCTTGTTCTGTCGTATCTCTTCCTCAGCTTCCTGAACGAGTCATGCACGATTAGACAGTTCTCTAGGTTCATTCCTCCCCTCATAATCATCACAAACGTGTCGCTTTTCCTCTCGGCAACGGTGGCAGGAGCCTTTTTCAAGCTCCGGGAGAAGTTAGCATTCCAGCAGAATCAGGTCCTGCTCTCGGGAATCTTCATCTTCCAGGGCTTTCTGGTGGTCCTGGTGATCTTCTTAAAGATATACCTCGAAAGAGTGACAATGAAGAGGCCCCTGTTCATCGTCTCCTCGGGGTCAAGGAGAAAGAAGGCCAAGGCCAACGTGTCCTTTTCAGAAGGCCTTGAGATCATGTCCCAGTCAAAGCTGCTGCTTGCAATGTCCCTCATCGTCCTCTTCTTCAACATCTCCTACAACATGGTCGAGTCCACATTCAAAGTCGGAGTTAAGGTTGCTGCAGAATACTTCAACGAGGAAAAGGGGAAATACTCCGGAAAGTTCAACCGGATCGACCAGTATATGACCTCTGTCGTGGTCATATGCCTCAACCTGTCACCATTCTCGAGCTATGTCGAGACAAGAGGGTTTCTTCTGGTCGGGCTTATAACGCCTATTGTGACACTCATGGCCATCGTTCTTTTCCTAGGGTCTGCGCTTTACAACACTAGCATGGAAGAGTCTGGGCTTGGAATCGTGAACGGGCTGTTTCCAGGTGGAAAGCCCCTGTACGTCCTCGAGAACTACTTCGGCGTAATCTTCATGTCGCTTCTGAAGATCACGAAGTACTCTGCATTTGACATCTGCAAAGAAAAGCTTGGGATGCGGATCAACCCCACATACAGAGCAAGGTTCAAGAGCGTGTATGATGGGATTTTTGGGAAGCTTGGGAAGTCCATAGGATCGATATACGGGCTCCTGATGTTCGAGGCCTTGGACACGGAAGACCTTAGAAAGGCAACGCCGATAACAGCAGGGATTATCTTTATCTTCATTGTCATGTGGGTAAAAGCGATAATCTACCTGTCGAGGTCTTACGAGTCTGCTGTACAGCATAATAGAGATGTTGACATCGACATGACCGAAAAGGCAAAGAAAAGCTTGGAGACTCCTGAGGAGCCAAAAGTTGTAGAT*GCAGGTGCTGGTGCTGGTGCTGGAGCAATTCTGTCTAAAGGTGAAGAATTATTCACTGGTGTTGTCCCAATTTTGGTTGAATTAGATGGTGATGTTAATGGTCACAAATTTTCTGTCTCCGGTGAAGGTGAAGGTGATGCTACTTACGGTAAATTGACCTTAAAATTTATTTGTACTACTGGTAAATTGCCAGTTCCATGGCCAACCTTAGTCACTACTTTCGGTTATGGTGTTCAATGTTTTGCGAGATACCCAGATCATATGAAACAACATGACTTTTTCAAGTCTGCCATGCCAGAAGGTTATGTTCAAGAAAGAACTATTTTTTTCAAAGATGACGGTAACTACAAGACCAGAGCTGAAGTCAAGTTTGAAGGTGATACCTTAGTTAATAGAATCGAATTAAAAGGTATTGATTTTAAAGAAGATGGTAACATTTTAGGTCACAAATTGGAATACAACTATAACTCTCACAATGTTTACATCATGGCTGACAAACAAAAGAATGGTATCAAAGTTAACTTCAAAATTAGACACAACATTGAAGATGGTTCTGTTCAATTAGCTGACCATTATCAACAAAATACTCCAATTGGTGATGGTCCAGTCTTGTTACCAGACAACCATTACTTATCCACTCAATCTGCCTTATCCAAAGATCCAAACGAAAAGACAGACCACATGGTCTTGTTAGAATTTGTTACTGCTGCTGGTATTACCCATGGTATGGATGAATTGTACAAATAA*

| **Primer** | **Sequence** |
| --- | --- |
| **PH-F** | ATGAGGACATTCAAAGAAATCGTTTACAA |
| **PH-R** | CTTGTCTTGTATATTCGAAGTAGATGTTAA |
| **NTT1-F** | TTAACATCTACTTCGAATATACAAGACAAG AATGAGGTGGAAAACAACAACCACAGTTTT |
| **NTT1-R** | ATCTACAACTTTTGGCTCCTCAGGAGTCTC AATTGCTCCAGCACCAGCACCAGCACCTGC |
| **GFP-F** | GCAGGTGCTGGTGCTGGTGCTGGAGCAATTCT |
| **GFP-R** | TTATTTGTACAATTCATCCATACCATGGGT |
| **attB1-F** | GGGGACAAGTTTGTACAAAAAAGCAGGCTATGAGGACATTCAAAGAAATCGTTTACAA |
| **attB2-R** | GGGGACCACTTTGTACAAGAAAGCTGGGTCTTATTTGTATAGTTCATCCATGCCATG |
| **attb_NTT1_R** | GGGGACCACTTTGTACAAGAAAGCTGGGTCTTAATCTACAACTTTTGGCTCCTCAGGAGT |
